# Local chromatin, 3D genome folding and transposon cargo shape Sleeping Beauty transposition

**DOI:** 10.64898/2026.11.03.763895

**Authors:** Christina J.I. Moene, Mathias Eder, Lise Dauban, J. Omar Yáñez-Cuna, Acadia A. Kocher, Laurens Eschauzier, Martijn Verkuilen, Vinícius H. Franceschini-Santos, Bas van Steensel

**Author notes:** These authors contributed equally to this work (Christina J.I. Moene, Mathias Eder).

## Abstract

The cut-and-paste DNA transposon Sleeping Beauty (SB) is widely used for genome engineering, mutagenesis and gene delivery. When mobilized from a genomic donor position (“launch site”), SB preferentially reintegrates locally (< 2 Mb), but factors governing its transposition from genomic sites remain incompletely understood. Here we analysed local transposition experiments in mouse embryonic stem cells to determine the influence of chromatin context and transposon cargo on SB transposition. Transposons in launch sites with transcriptionally active, accessible chromatin showed lower transposition efficiency. Large cargo and cargo with higher transcriptional activity also result in reduced transposition efficiency. After excision, local reintegration patterns closely resembled 3D contact frequencies of the launch site. This indicates that chromosome folding is a major determinant of the transposition pattern. Surprisingly, we did not detect any effect of the destination chromatin composition on reintegration efficiency. Cargo sequence modified the reintegration patterns: promoter-containing cargo increased both very proximal (<10kb) and interchromosomal transpositions, while CTCF binding sites in the cargo increased reintegration on the launch chromosome. Together, these results define key determinants of SB transposition in chromosomes and inform the design of SB-based genome engineering strategies.

## Introduction

The DNA transposon Sleeping Beauty (SB) is a cut-and-paste transposon capable of excising a DNA element from one genomic locus and reintegrating it into another. SB is a reconstructed member of the Tc1/mariner superfamily that was restored from inactive fish transposon sequences and shown to transpose in vertebrate cells (*1*). SB transposition requires two elements that can be supplied separately: the SB transposase and the two terminal inverted repeat/direct repeat (IR/DR) sequences of the transposon (*2*). During transposition, the transposase binds the transposon ends, excises the element from its donor site, and inserts it into a new genomic location. Excision leaves a characteristic CAG footprint at the donor site, whereas reintegration occurs predominantly at TA dinucleotides (*3*).

The SB transposon system can be used to integrate exogenous sequences (“cargo”) into a mammalian genome, from a donor plasmid containing the SB transposon with the cargo (*1, 4*). This has applications in the generation of transgenic cell lines, generation of germline-transgenic animals and gene delivery in gene therapy (as reviewed in (*5*)).). In addition, SB transposase can mobilize and reintegrate a genomically integrated SB transposon (*3*). We refer to such a defined genomic donor position as the “launch site”. When mobilized from the genome, SB has a preference for local reintegration (*3, 6, 7*). Therefore, re-mobilization of a genomic SB transposon has been used for detailed exploration of genomic loci, using insertional mutagenesis (*6, 8–11*) or insertion of reporter genes (*12, 13*), both in cell culture and *in vivo* (*11*). This genomic transposition is the main focus of this article.

Earlier work showed that SB transposition is influenced by both transposon design and launch site context. The activity of the system was greatly improved by engineering hyperactive transposase variants, most notably SB100X (*14*) which is used throughout this study. In addition, transposition can be enhanced when the transposon is CpG methylated or embedded in heterochromatin, indicating that chromatin context at the launch site can affect transposition efficiency (*14*). Excision and reintegration can also be mechanistically separated, with engineered SB variants that remain excision proficient but are defective in reintegration (*15*). Cargo size has previously been reported to reduce SB transposition efficiency when mobilized from a plasmid (*16*), but the effect of cargo on excision from the genome has to our knowledge not been reported.

Compared with retroviral and lentiviral vectors, SB shows a relatively weak bias towards integration in active genes and has therefore become an attractive tool for stable gene delivery (*14*). At the sequence level, however, SB integration is not random. In addition to its strong requirement for a TA dinucleotide, recent work identified a broader local target-site preference centred on an ANNTANNT motif (*17*).

Despite these insights, it is still not well understood how strongly the genomic launch position of the transposon influences transposition efficiency of a genomically integrated SB transposon. It is also unclear to what extent properties of the transposon cargo itself, such as size or transcriptional activity, affect excision and reintegration. Finally, although the bias for local reintegration has long been recognized, the relationship between local transposition patterns and higher-order genome organisation has not been examined systematically. Knowledge of the properties of SB transposition from the genome would improve the design of studies using this system.

To address these questions, we used a collection of mouse embryonic stem cell lines carrying either single-copy or multiple SB transposons at defined genomic positions and containing different cargos. By combining previously generated datasets (*13, 18, 19*) with new transposition experiments, we examined how launch site chromatin context, 3D genome folding and cargo properties influence transposition. First we examine how these factors affect the efficiency of transposition, then we explore how they determine where SB reintegrates, both locally and genome-wide.

## Results

### Differences in transposition due to genomic location

To study the characteristics of SB transposition within the genome of mouse embryonic stem cells, we initially used previously generated clonal cell lines that each carry a single SB transposon at a distinct genomic location (**Fig 1A, Table S1**). Each transposon contains the same 335 bp cargo: a loxP site and flanking sequences, which is not expected to have any transcriptional or regulatory activity (*18*). Throughout this manuscript, we define transposition as excision of the transposon from the launch site followed by reintegration elsewhere in the genome. The genomic context surrounding each launch site is shown in **Fig S1**.

**Figure 1.**
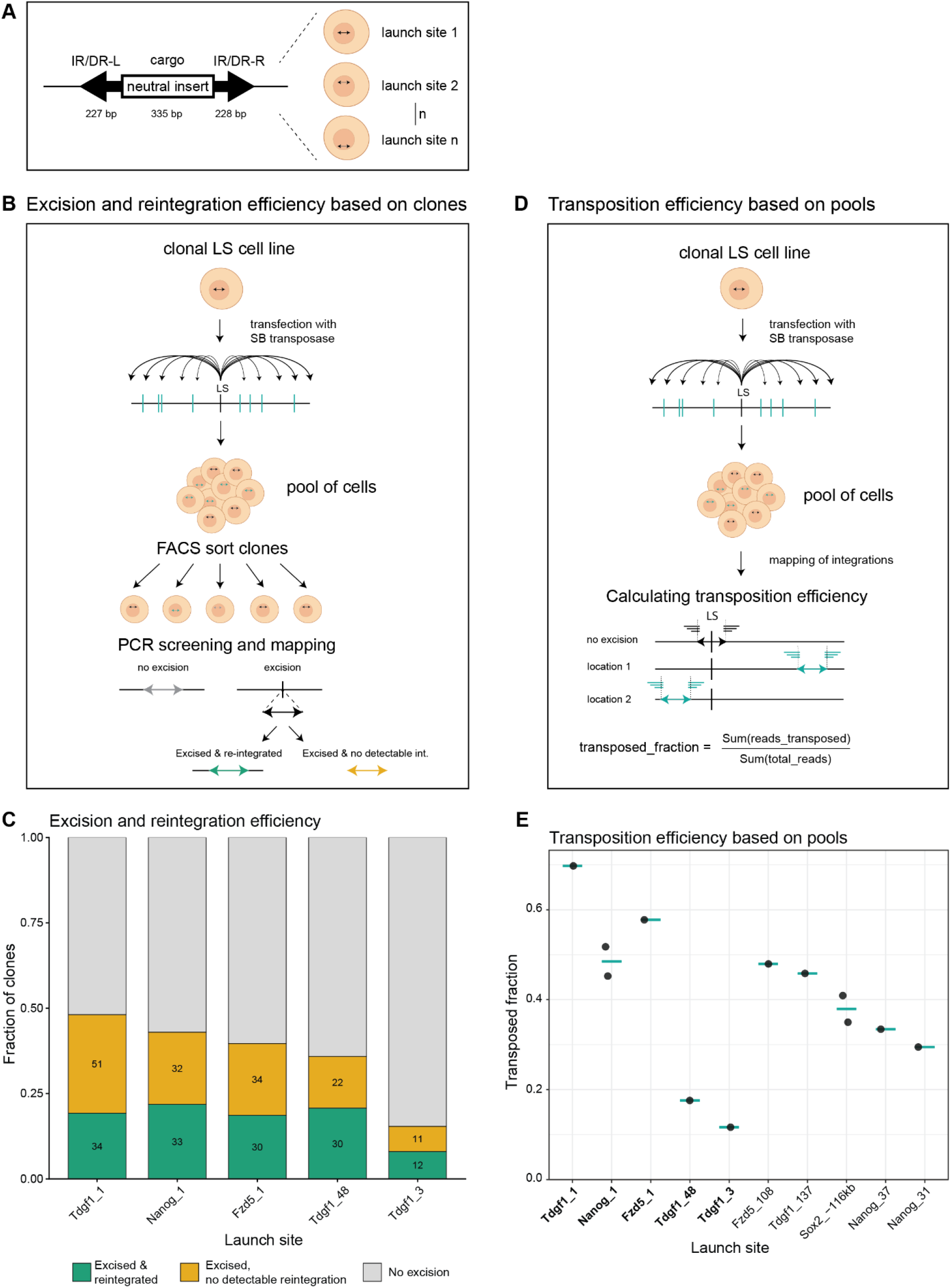
Assessing transposition efficiencies based on clonal and pool-based data. **(A)** Schematic of the Sleeping Beauty (SB) transposon used in this study. A neutral DNA cargo is flanked by the terminal inverted repeat/direct repeat sequences (IR/DRs). Each cell line carries a single transposon integration at a distinct genomic locus, here referred to as the launch site. **(B)** Schematic workflow for the clone-based assay used to classify transposition outcomes after SB transposase expression. Clones were scored as no excision, excision with detectable reintegration or excision without detectable reintegration. **(C)** Fraction of clones per launch site in each outcome class: excision with reintegration (green), excision without detectable reintegration (yellow) or no excision (grey). Numbers indicate the number of clones for which a reintegration was or was not found in the mapping. **(D)** Schematic workflow for the pool-based assay to estimate transposition efficiency. The transposed fraction was calculated as the number of reads derived from transposed reintegrations divided by the total number of reads. **(E)** Transposed fraction in pool-based transposition assays, mobilizing SB with a neutral cargo from different genomic launch sites, calculated as shown in (D). Dots represent biological replicates of the transposition experiment, green lines indicate the mean. Launch sites also assessed in C are marked in bold. LS: launch site.

We first asked whether the same transposon shows different transposition efficiencies when launched from different genomic loci. We transfected five launch site cell lines with SB transposase and isolated single clones after six days. We screened these clones by PCR across the launch site to identify excision events (**Fig S2**), as described previously (*18*), and then mapped transposon insertions using a Tn5-based sequencing approach (*20*). This workflow allowed us to distinguish clones with no excision, clones with excision and detectable reintegration, and clones with excision but no detectable reintegration (**Fig 1B**).

Clonal analysis revealed marked differences between launch sites (**Fig 1C, Table S2**). The fraction of excision-positive clones varied from 15% for launch site Tdgf1_3 to 48% for Tdgf1_1. By contrast, among excision-positive clones, the proportion with a detectable reintegration event was more consistent (40% – 52%). These results suggest that the genomic location of the launch site has a stronger effect on excision than on reintegration after excision.

We then assessed transposition efficiency in ten launch site cell lines, including the five previously studied. To streamline this analysis, we directly analysed the pooled cell populations generated after transposase expression (**Fig 1D**). We mapped integrations in these bulk populations and calculated for each cell line the transposed fraction as the number of reads derived from transposed integrations divided by the total number of mapping reads. This analysis again showed large differences between launch sites, with transposed fractions ranging from 12% for Tdgf1_3 to 70% for Tdgf1_1 (**Fig 1E**).

For the five launch sites analysed by both approaches, the clone-and pool-based measurements show a qualitatively similar trend (e.g., Tdgf1_3 has the lowest transposition efficiency), but the pool-based estimates of transposition frequency are higher, even when correcting for excised-not-reintegrated transposons which are not detectable in the pool (not shown). This might be the result of differences between the two assays. In the pool-based assay off-target reads will be counted among the ‘transposed integrations’, while the clone-based assay has stricter filtering and might instead miss integrations in poorly-mappable regions. In addition, differences in PCR-efficiency between the launch site and reintegration sites could bias the pool-based estimates. Finally, clones that retain the transposon at the launch site while also acquiring an additional insertion, for example due to transposition after DNA replication, are marked as ‘no excision’ in the clone-based assay but will contribute to the transposed fraction in the pool-based assay. However, overall these results show that genomic position affects SB transposition efficiency, primarily through modulating the mobilization efficiency. This motivated us to examine which genomic features underlie this effect.

### Active chromatin launch site reduces transposition efficiency

To systematically test whether the chromatin context of the launch site influences SB transposition, we measured transposition from multiple SB launch sites located in diverse chromatin locations in clonal cell lines. For this, we generated a nested transposon construct in which a SB transposon carrying a short neutral cargo (239 bp, loxP site) was embedded within a PiggyBac (PB) transposon (**Fig 2A**). Co-transfection of this plasmid construct with PB transposase generated clonal cell lines carrying multiple chromosomal SB launch sites. We isolated individual clones, determined the genomic positions of the PB insertions by sequencing outwards from the PB transposon arms, and selected three clones (named WT78, WT79 and WT83) carrying 14 – 19 nested PB-SB insertions each. In this system, all SB launch sites have exactly the same immediate SB-flanking PB sequences and reside within the same clonal background, minimizing variation caused by launch site sequence differences, clonal variation or transfection efficiency. This design therefore allows us to directly assess the effect of genomic location, and thus chromatin context, on SB transposition.

**Figure 2.**
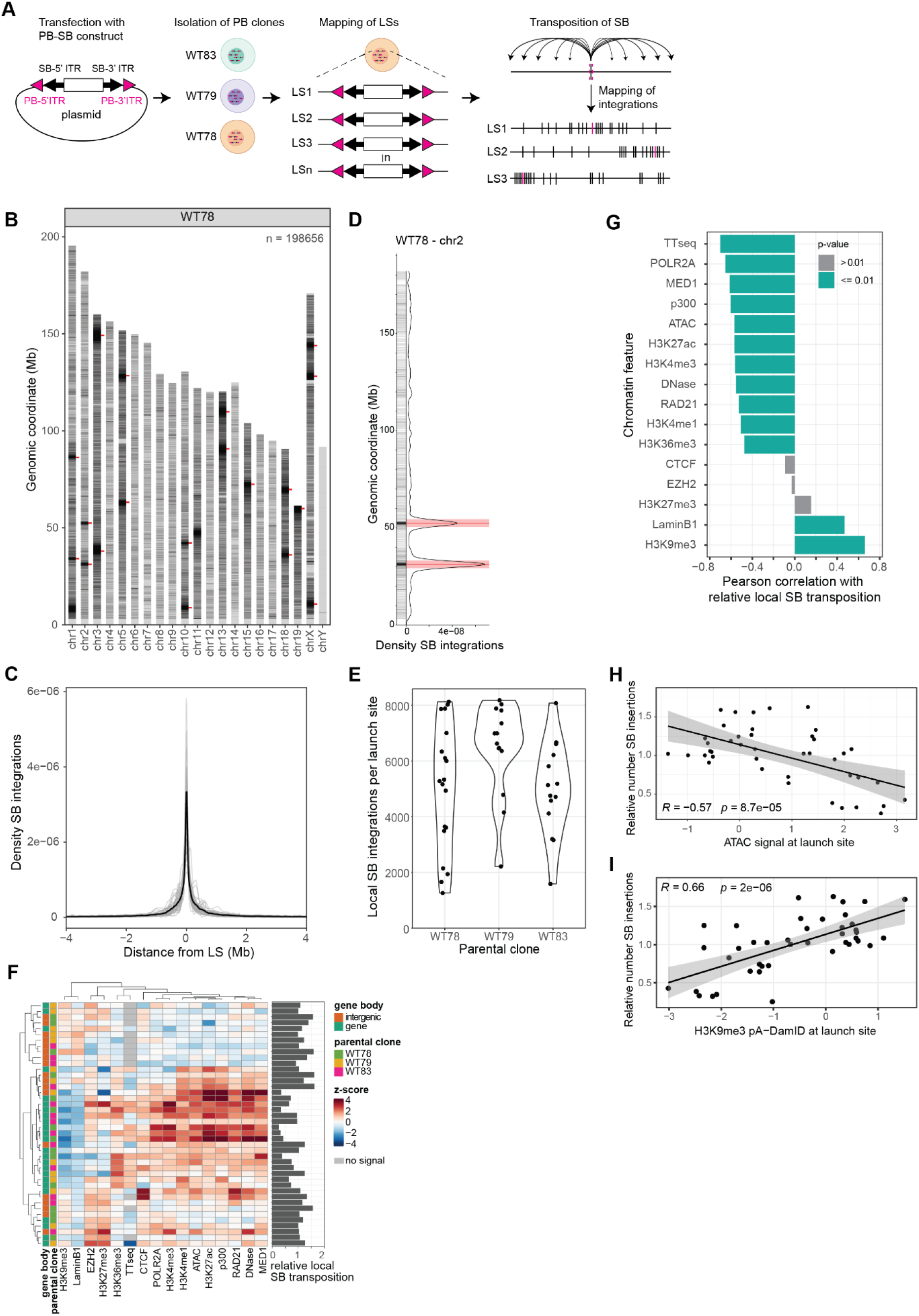
Chromatin context launch site affects transposition efficiency. **(A)** Experimental set-up: cells are transfected with a nested transposon construct; clonal cell lines are isolated; PB integrations are mapped in each clone by sequencing; SB transposition is induced by SB transposase transfection and reintegrations are mapped by sequencing. LS: launch site. **(B)** SB reintegrations mapped after SB transposase transfection in WT78. Red ticks indicate the mapped launch sites. Two biological replicates combined. n = total number of mapped SB reintegrations. **(C)** Density of SB reintegrations around the launch sites, per launch site (light grey) and for all launch sites combined (black). **(D)** SB reintegrations mapped in WT78 chromosome 2, individual reintegrations and density plot. Red lines indicate launch sites, red shaded areas indicate the ±2Mb region used for quantification of local SB reintegrations. **(E)** Number of SB reintegrations mapped within 2Mb of each launch site, per parental clone. **(F)** Heatmap of the levels of 16 chromatin features in a 2kb window around each of the 42 launch sites. Features are z-score normalized based on a genome-wide background set, for visualization the scale is truncated at [-4,4]. ChIP and pA-DamID data are pseudocounted, for TTseq no signal is indicated in grey. Source data provided in Methods. Side barplot indicates the relative local SB transposition per launch site (number of local reintegrations, normalized to the mean per parental clone). **(G)** Pearson correlations between relative local SB transposition and the signal for each of the 16 chromatin features (ordered by correlation coefficient) across the 42 launch sites. Color indicates significance (p ≤ 0.01). For TT-seq, launch sites with no signal are excluded. **(H-I)** Example plots showing the relationship of relative local SB transposition with ATAC-seq (**H**) or H3K9me3 pA-DamID signal (**I**) at the launch site, across the 42 launch sites. R is Pearson correlation coefficient. **(C-I)** Two biological replicates for three parental clones combined. **(G-I)** Chromatin features are calculated as in F, but not truncated.

After induction of SB transposition in the three parental clones in two biological replicates, we mapped more than 500,000 SB reintegrations in the resulting pools (**Fig 2B, S3A**). Across the three parental clones, 52-57% of mapped reads correspond to the launch sites (PB sequence), indicating that of the detectable SB transposons 45% had mobilized and reintegrated (mobilized and not-reintegrated transposons are undetectable in this experiment). Of the ones that transposed, 47% of reintegrations were located within 2 Mb of a mapped launch site (**Fig 2C**), indicating strong local enrichment. The remainder of reintegrations were detected throughout the genome, including on chromosomes without a mapped launch site. A few distal reintegration hotspots without a nearby mapped launch site (e.g. WT78, chr1, around 9 Mb) likely originated from launch sites located in poorly mappable regions.

Because all SB launch sites have an identical sequence, reads from non-mobilized SB cannot be assigned to a specific position. Therefore, we cannot compute the transposed fraction per launch site. Instead, we therefore estimated launch site specific local transposition efficiency by counting SB reintegrations within ±2 Mb of each launch site (**Fig 2D**). This yielded 1260-8178 reintegrations per launch site (**Fig 2E**), of which we expect fewer than 75 to be derived from a different launch site (based on the density of reintegrations on chromosomes without a launch site). There is up to 6-fold variation in the number of mapped reintegrations per launch site within the same parental clone (**Fig 2E**). After internal normalization per clone, the relative local SB transposition was highly reproducible between biological replicates (**Fig S3B**), indicating that these differences reflect properties of the launch sites rather than stochastic variation in the experiment.

We characterized the chromatin context of each launch site by quantifying the abundance of 16 chromatin marks in a 2-kb window around the launch site, using publicly available epigenome data (**Table S3**). Because PB insertions generally acquire the chromatin properties of their genomic environment (*21–23*), the chromatin state in wild-type cells should also describe the launch sites. After restricting the analysis to autosomal launch sites in well-mappable regions, 42 launch sites remained. These loci spanned a broad range of chromatin environments, including lamina-associated domains and various heterochromatic and euchromatic features (**Fig 2F**). Consistent with the known preference of PB insertions for accessible chromatin (*24*), these launch sites were modestly enriched for euchromatic features (**Fig S3C**).

To assess the effect of chromatin context on local transposition frequency, we computed the Spearman correlations between the relative number of SB reintegrations around the 42 launch sites and each chromatin feature. Overall, the number of reintegrations is negatively correlated with various features of active and accessible chromatin and positively correlated with heterochromatin marked by H3K9me3 and LaminB1 (**Fig 2G**), regardless of the window size used for the chromatin context quantification (**Fig S3D**). For many features, such as H3K9me3 and chromatin accessibility (measured by ATAC [Assay for Transposase-Accessible Chromatin]), changes in the feature directly relate to a change in transposition efficiency across the whole range covered by our launch sites (**Fig 2H-I, S3E**). Polycomb-related heterochromatin (marked by H3K27me3 and EZH2) does not correlate with transposition efficiency (**Fig 2G**). Note that many of these chromatin features are mutually correlated across the genome, so the relationships with transposition efficiency are not independent.

RNA polymerase II (POLR2A) ChIP-seq and nascent transcription (TTseq) are among the features most strongly anticorrelated with transposition efficiency (**Fig 2G, S3E-F**). We wondered whether this was the result of poor transposition from sites close to active cis-regulatory elements (i.e., elements that are transcriptionally active and enriched in active chromatin features), or that transcription through the launch site might impair transposition independent of other active marks. To test this, we grouped the launch sites by the presence or absence of a proximal putative cis-regulatory element (here defined by the presence of a DNase-hypersensitivity peak in the 4kb window around the launch site) (**Fig S3G**). We found that the effect of RNA polymerase II and TTseq on transposition efficiency is strongest for launch sites with a proximal DNase peak, but that there is also a negative correlation among launch sites without a proximal cis-regulatory element. This indicates that transcription itself might impair transposition (**Fig S3G**).

Finally, we asked whether the variation in transposition efficiency observed across cell lines carrying a single SB launch site (**Fig 1D**), could also be explained by differences in chromatin context. These 10 launch sites are located in different chromatin contexts, but the variation is smaller than between the 42 launch sites created with the nested PB-SB transposon (**Fig S4A**). Here, the transposed fraction per launch site does not correlate significantly with any of the tested chromatin marks (**Fig S4B**). This likely reflects a combination of reduced chromatin variation, greater experimental variability between independently transfected cell lines, differences between launch sites in surrounding sequence context and the small number of tested launch sites. Together, these results indicate that launch site chromatin can have a strong effect on SB transposition efficiency, but additional factors may play a role.

### Transposition efficiency depends on cargo activity and size

We next asked whether properties of the SB cargo influence transposition efficiency. To address this question, we compiled datasets from transposition pools derived from single launch site cell lines carrying cargos with distinct sequence features. These cargos included neutral sequences, genomic promoters, synthetic promoters, a gene, an enhancer and one or multiple CTCF-binding sites (CBSs) (**Fig 3A**). Overall, the cargos varied in their transcriptional activity, binding of transcription factors, and in size. To reduce confounding effects by genomic position, we first compared different cargos mobilized from the same launch site (three sites: Sox2_-161kb, Sox2_-116kb and Nanog_1). We quantified transposition efficiency as the transposed fraction, calculated as described in **Fig 1D**.

**Figure 3.**
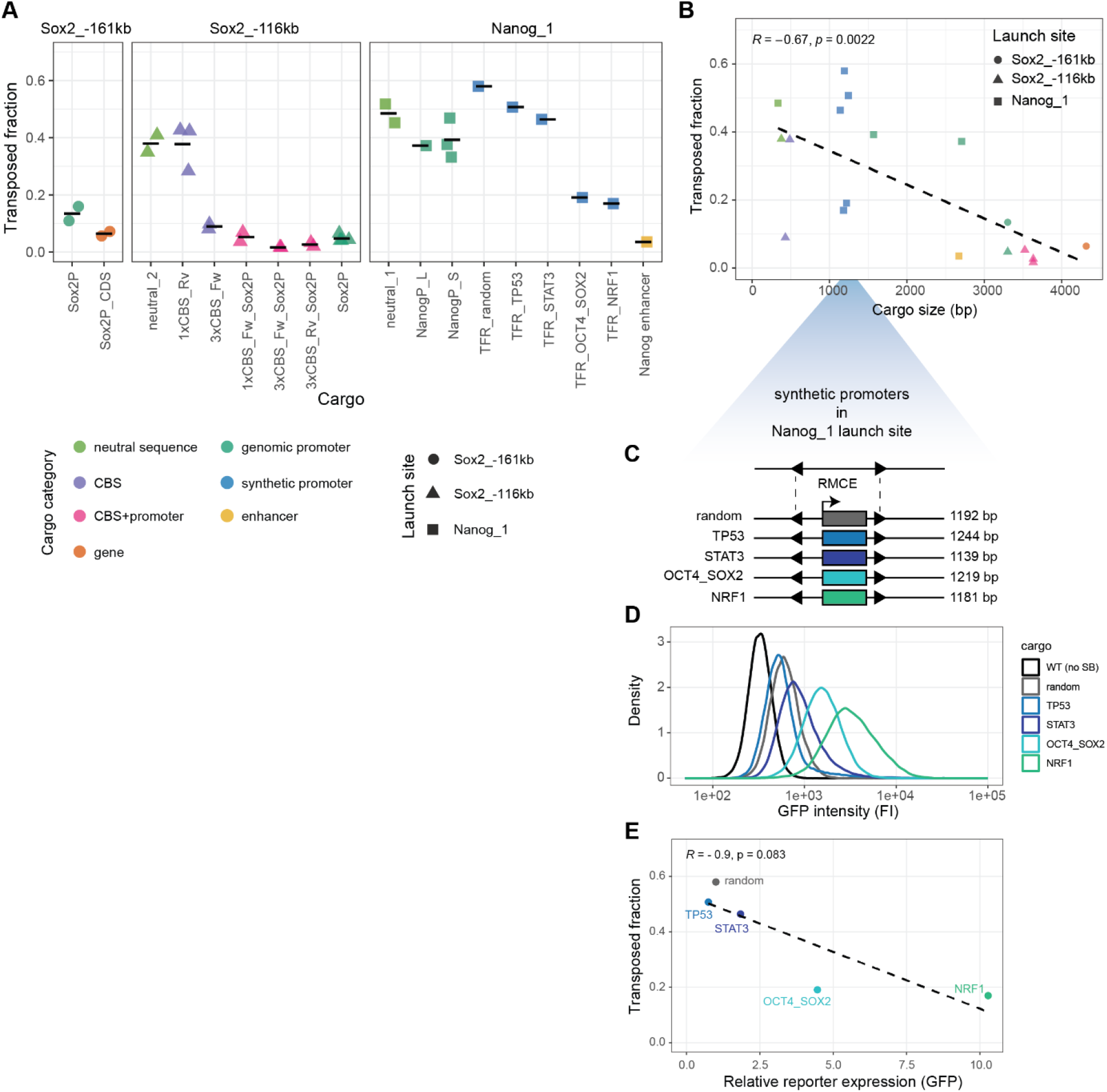
Transposition efficiency depends on cargo activity and size. **(A)** Transposed fraction in pool-based transposition assays, for different transposon cargos at the same genomic launch sites (Sox2_-161kb, Sox2_-116kb and Nanog_1). Shapes indicate launch site, colours cargo category, dots represent biological replicates, and horizontal lines represent the mean per cargo. TFR is transcription factor reporter, CDS is coding sequence, NanogP is Nanog promoter (L long, S short), Sox2P is Sox2 promoter. **(B)** Relationship between cargo size and mean transposed fraction, from the data in A. Shapes indicate launch site, colors indicate cargo category. Biological replicates of the same cargo are averaged. R is Spearman correlation coefficient. **(A-B)** For NanogP_S three similar constructs (sizes 1485-1671 bp) are treated as biological replicates, using the mean size. **(C)** Schematic of the different synthetic promoter cargos in launch site Nanog_1. Each construct contains four identical transcription factor binding sites driving GFP expression. RMCE: Recombination-Mediated Cassette Exchange.**(D)** Activity of the synthetic promoter cargos in the same launch site measured as GFP expression by flow cytometry. Colour indicates the synthetic promoter identity. **(E)** Relationship between relative reporter expression of the synthetic promoters and their transposed fraction. Relative reporter (GFP) expression is calculated by subtracting autofluorescence (based on a wild-type control) and normalizing to the random reporter. R is Spearman correlation coefficient, colour indicates the synthetic promoter identity, one biological replicate.

Across all three launch sites, different cargos showed a broad range of transposed fractions (**Fig 3A**). The short neutral sequences generally had a high transposed fraction (38% and 49%), whereas several promoter- or CBS-containing constructs had a lower transposition efficiency. In Sox2_-116kb, constructs that combine CBSs with the *Sox2* promoter were among the least efficient (transposed fraction 2-5%). These comparisons indicate that the cargo sequence contributes to transposition efficiency in addition to the genomic context of the launch site.

Across all launch site backgrounds, larger cargos tended to transpose less efficiently than smaller ones (Spearman R = −0.66, p = 0.0031; **Fig 3B**). However, compared to the small neutral sequence, we also expect the largest cargos (promoters and an enhancer) to be bound by more transcription factors and be more transcriptionally active. Therefore, this correlation might be driven by a combination of cargo size and cargo activity. To control for this, we compared two *Nanog* promoter reporters mobilized from the SB launch site. The NanogP_L construct is 1036 bp larger than NanogP_S (2707 bp versus 1671 bp) and showed a lower transposed fraction, despite having similar reporter expression (**Fig S5A**). This comparison supports the conclusion that insert size itself contributes to reduced transposition efficiency, which is in line with the effect of cargo size on SB mobilization from a plasmid (*19*).

Cargo size alone is unlikely to explain all variation, because several synthetic promoters mobilized from launch site Nanog_1 had similar sizes but different transposition efficiencies (**Fig 3B**). These synthetic promoter reporters each consist of four identical transcription factor binding sites and a minimal promoter driving GFP expression (*25*); a construct with four random sequences (devoid of transcription factor binding motifs) serves as an inactive control (**Fig 3C**). Flow-cytometry measurements showed an approximately 10-fold range in GFP expression across these reporters (**Fig 3D**). Higher reporter activity was associated with lower transposed fractions (**Fig 3E**), although this trend did not reach statistical significance in this small set (R = −0.9, p = 0.083). These data therefore suggest that cargo with higher transcriptional activity reduces transposition efficiency.

To further examine the relationship between reporter activity and transposition efficiency, we re-analysed datasets generated in previous *Sox2* reporter studies (*13, 19*). First, we used data where the activity of a *Sox2* promoter reporter (Sox2P) was reduced due to CBS deletions in and around its nearby enhancer, the Sox2-control region (SCR) (*19*). Although these CBS deletions reduced Sox2P reporter expression, they did not detectably alter transposition efficiency from the Sox2_-116kb launch site (**Fig S5B**). However, in this setting both reporter expression and transposition efficiency were already low at baseline, which may have limited our ability to detect a further reduction in transposition.

Second, we re-analysed data from *Sox2* gene deletion experiments in which two different reporter cargos were mobilized from the same Sox2_-161kb launch site (*13*). These consisted of the Sox2P reporter and a related reporter that additionally contains the *Sox2* coding sequence (Sox2P_CDS). The endogenous *Sox2* gene (52 kb downstream of the launch site) was deleted, which increased expression of both reporters, although to different degrees, and this increase coincided with reduced transposition frequencies (**Fig S5C**). Together, these comparisons support a negative relationship between transcriptional cargo activity and transposition efficiency, but they also indicate that this effect is context-dependent.

Overall, these results show that transposition efficiency depends not only on the chromatin environment of the launch site but also on intrinsic properties of the transposon cargo, where larger and more active cargos tend to transpose less efficiently.

### Local reintegration pattern follows 3D contact with the launch site

After assessing the factors influencing SB transposition efficiency, we turned to analyse the locations where SB reintegrates. Previous work showed that SB integrates preferentially at TA dinucleotides (*3*), with a milder preference for the motif ANNTANNT (*17*). Across more than 500,000 mapped reintegrations, we found that 95.5% occurred at TA sites. In addition, we also recovered the two extra preferred bases at the fourth nucleotide upstream and downstream of the insertion site (**Fig 4A**). Each of the flanking bases occurred at 65% of the reintegration sites, and 93% of reintegrations contained at least one of them (**Fig S6A**). Both extra bases and the full motif were enriched over their abundance in the genome (**Fig S6B**). Finally, we checked whether there is a secondary preferred sequence among the reintegrations not at a TA site (4.5% of reintegrations), but those do not seem to have any discernible preferred motif (**Fig S6B**, note the y-axis scale). These results confirm the canonical TA preference of SB and further refine the local sequence context preferred during reintegration.

**Figure 4.**
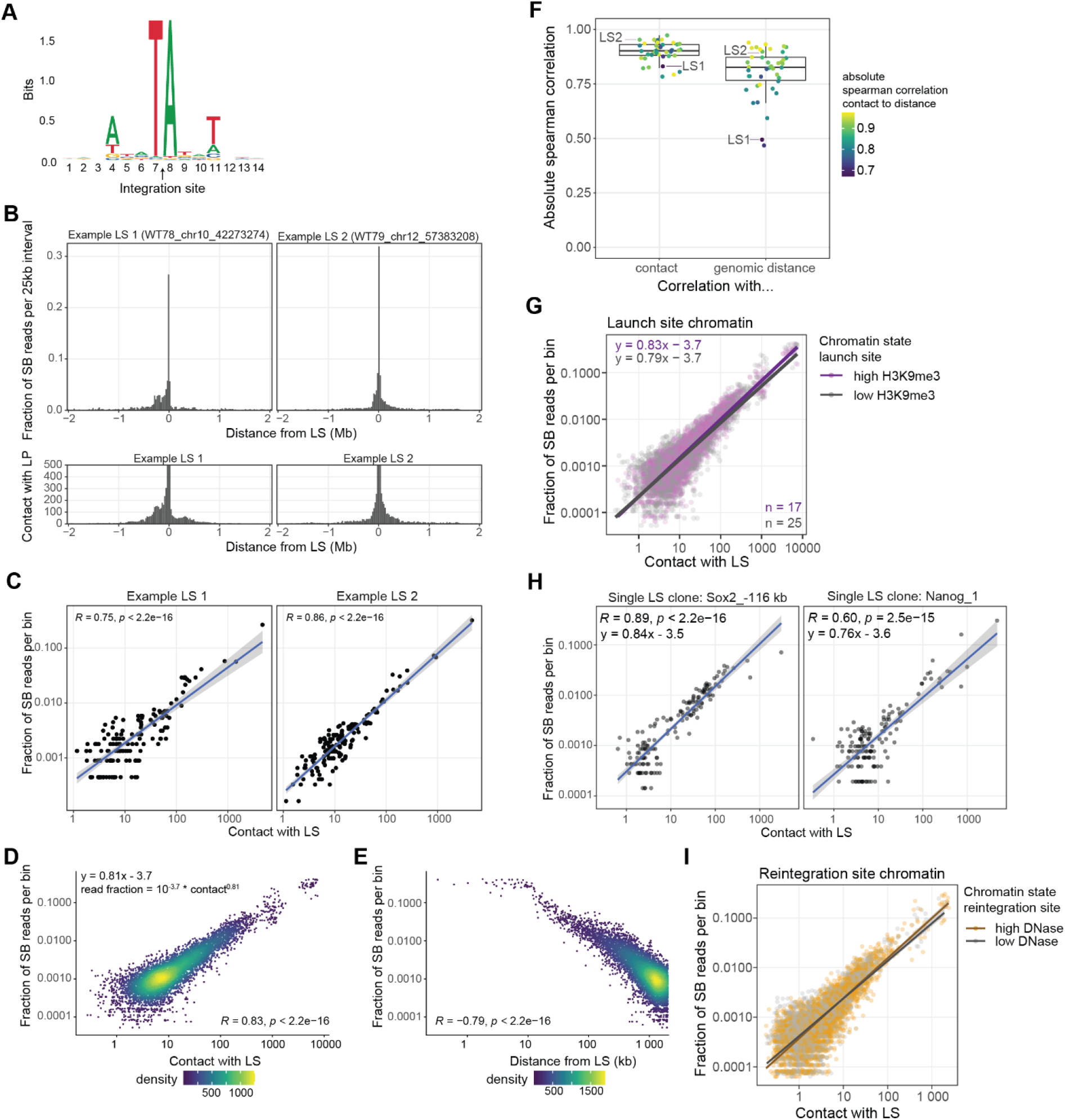
3D-contact determines reintegration distribution. **(A)** Sequence motif of the SB reintegration sites, based on the data described in Fig 2. **(B)** For two example launch sites from nested transposon clones WT78 and WT79, Top: distribution of SB mapping reads (proxy for reintegration events) within ± 2Mb of the launch site. Bottom: Virtual 4C contact profile (from micro-C data, (*26*)) with the location of the launch site as the viewpoint. Top and bottom are calculated on the same 25 kb bins. **(C)** Correlation between the fraction of SB mapping reads and the contact with the launch site, for the same example launch sites as in B. **(D)** As in C, but for all 42 launch sites combined, and displayed as a density scatterplot. **(E)** As D, and using the same 25 kb bins as D, but for the relationship between the genomic distance from the launch site and the fraction of SB mapping reads. **(C-E)** R is Spearman correlation coefficient. **(F)** The absolute spearman correlation between SB mapping reads and 3D contact or genomic distance for each of the 42 launch sites. Color indicates the spearman correlation between 3D contact and linear distance for that launch site. Example launch sites shown in B-C are indicated. **(G)** Relationship between the fraction of SB mapping reads per bin and contact frequency of that bin with the launch site, stratified by high (z-score > 0) or low (z-score ≤ 0) H3K9me3 levels at the launch site. H3K9me3 level is calculated in a 2kb bin surrounding the launch site (as in Fig 2F). n indicates the number of launch sites in each group. **(H)** Relationship between the fraction of SB mapping reads and contact frequency in single launch site cell lines carrying a neutral transposon cargo at the Sox2_-116 kb or Nanog_1 launch site. Data from two biological replicates combined. R is spearman correlation coefficient. **(C-E, G-H)** Contact and SB mapping reads are calculated on 25 kb bins within 2 Mb of the launch site. Bins with 0 contact or reintegrations are excluded. **(I)** Relationship between the fraction of SB mapping reads per 10 kb bin within 2 Mb of a launch site and contact frequency of that bin with the launch site, stratified by high (z-score > 0) or low (z-score ≤ 0) DNAse accessibility at the 10 kb reintegration bin. Launch sites with less than 30% high or low bins are excluded. Bins with 0 contact or 0 reintegrations are excluded. **(A-G, I)** SB reintegration data from the nested transposon experiment described in Fig 2. Data pooled from three cell lines, two biological replicates per cell line.

After the base scale sequence preferences, we next analysed the local pattern of SB reintegrations around individual launch sites. Because reintegrations close to the launch site are strongly enriched, multiple independent transposition events may occur at the same TA site. In such regions, counting only unique genomic insertion sites can therefore underestimate the true number of local reintegrations. To reduce this bias, we used the distribution of unique mapping reads as a proxy for reintegration-event abundance in highly complex pools, as described in the Methods (‘Quantification of reintegration events’). We then quantified the fraction of mapping reads in 25 kb bins around each launch site in the nested-transposon cell lines. This gives a pattern of *where* SB reintegrates locally. Two representative launch sites illustrate that the local reintegration pattern can differ substantially in shape and symmetry (**Fig 4B**).

We wondered whether these patterns may be explained by the three-dimensional (3D) folding of the chromatin fibre. Indeed, the local reintegration profiles of these examples closely resembled the 3D contact profiles of the launch sites, as inferred from published Micro-C data (*26*) (**Fig 4B**). Consistent with this, the fraction of SB mapping reads per 25 kb bin correlated strongly with contact frequency for both example loci (**Fig 4C**). We observed the same pattern across all launch sites (**Fig 4D**). Thus, local reintegration patterns reflect locus-specific 3D genome folding.

As expected, the abundance of SB reintegrations per 25 kb bin also decreased with increasing linear genomic distance from the launch site (**Fig 4E**). However, within 500 kb of the launch site, where contact patterns are more variable, the correlation with contact (Spearman R = 0.88) (**Fig S6C**) was stronger than that with genomic distance (Spearman R = −0.78) (**Fig S6D**). Moreover, when analysed separately for each launch site, the Spearman rank correlation between SB reintegrations per bin and contact remained consistently high, whereas the correlation with linear genomic distance was weaker for a subset of launch sites (**Fig 4F**). The relationship between SB reintegrations per bin and contact was similar across launch sites with different chromatin states, including sites with high or low H3K9me3 signal (**Fig 4G, Fig S6E**). Finally, two independent datasets in which a single SB transposon carrying a short neutral cargo was mobilized from Sox2_-116kb or Nanog_1 showed the same overall behaviour (**Fig 4H**). Together, these analyses indicate that 3D contact is a better predictor of *where* local SB reintegration occurs than linear distance, especially in loci with more structured local folding.

### Reintegration site chromatin does not detectably affect reintegration efficiency

Finally, we asked whether the chromatin composition of the reintegration locations affects SB reintegration. We hypothesized that SB, once excised, might reintegrate more frequently in open chromatin. Since 3D contact is such a strong predictor of the reintegration pattern, we decided to test this in the context of the relationship between contact and reintegration. Therefore, we classified all 10 kb bins around the launch sites as ‘high’ or ‘low’ accessibility (as measured by DNase sequencing, **Table S3**) and restricted the analysis to launch sites surrounded by both classes of bins (at least 30% each). Across those launch sites, both high and low accessible regions have a very similar relationship between contact and SB reintegration (**Fig 4I**). In other words, regions with similar contact to the launch site but different accessibility will have a similar frequency of SB reintegration: the frequency of SB reintegration is independent of local chromatin accessibility.

We repeated this analysis for each of the 16 chromatin marks (**Table S3**) surveyed before. For none of these marks is there a detectable effect of the local level of the mark on the reintegration frequency at that location (**Fig S6F**). Thus, we found no evidence that local chromatin composition at the reintegration site affects the efficiency of SB reintegration.

### SB cargo influences local and distal reintegration patterns

So far, we have focused primarily on the transposition characteristics of short neutral cargos. However, we recently showed that adding CBSs to a promoter reporter can substantially alter where SB reintegrates (*19*). To further examine how cargo sequence influences SB reintegration location, we compared reintegration patterns between different cargos mobilized from the same launch sites.

First, we asked if the transposon cargo influences the overall transposition distribution. For this, we classified the detected reintegrations into three categories: local reintegrations within 2 Mb of the launch site, *cis*-distal reintegrations on the same chromosome but outside the 2 Mb window, and *trans* reintegrations on other chromosomes (**Fig 5A**).

**Figure 5.**
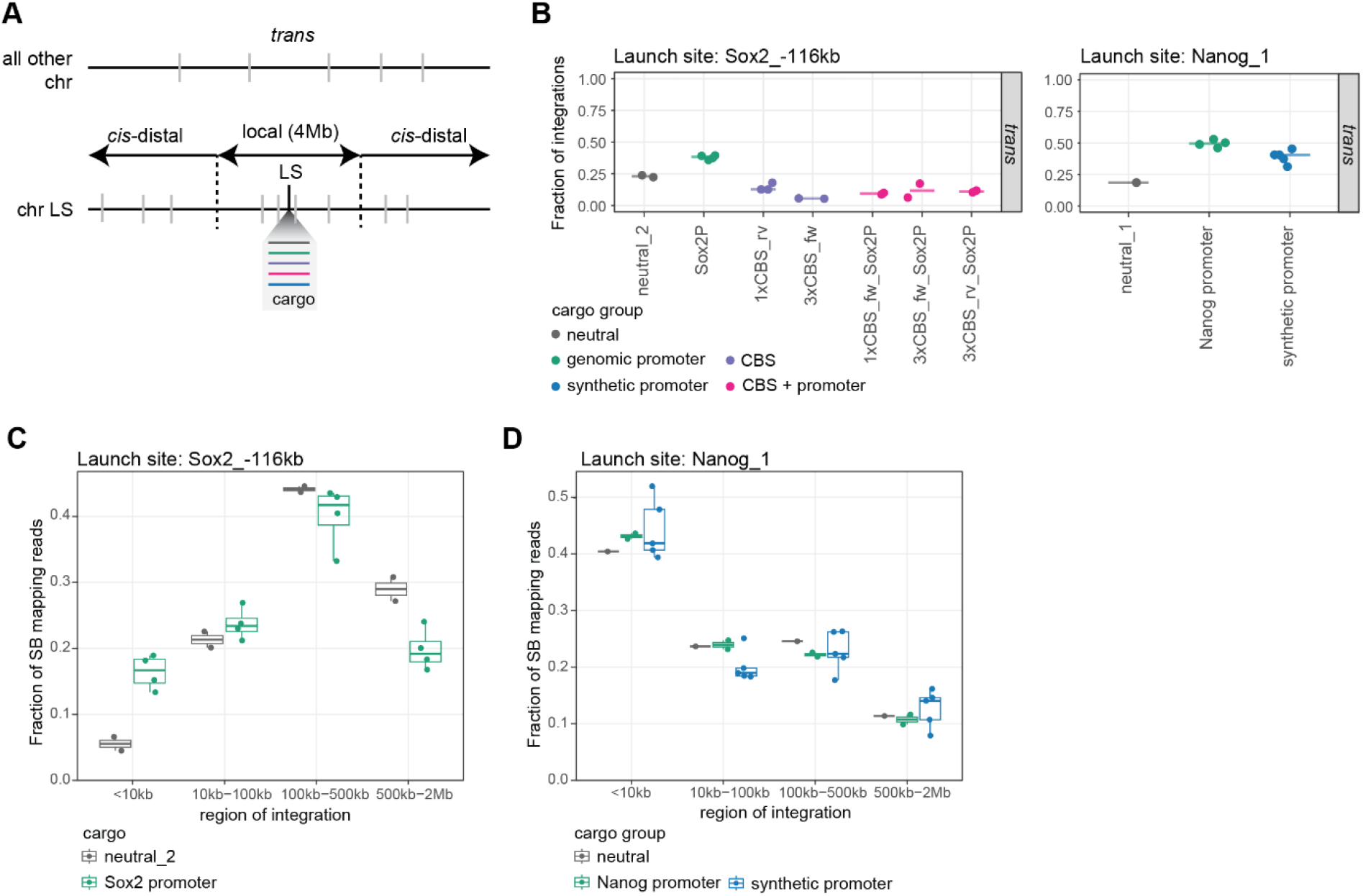
Cargo characteristics shape local and interchromosomal reintegration patterns. **(A)** Schematic representation of reintegration categories. Local reintegrations are defined as reintegrations within ±2 Mb of the launch site. *Cis*-distal reintegrations are located on the same chromosome but outside the ±2 Mb local window. *Trans* reintegrations are located on chromosomes other than the launch-site chromosome. Different colored lines indicate different transposon cargos mobilized from the same launch site. **(B)** Fraction of reintegrations classified as trans for different cargos mobilized from the Sox2_-116 kb and Nanog_1 launch sites. Dots represent biological replicates (Sox2_-116kb) or reporter constructs (Nanog_1) and horizontal lines indicate the median. **(C)** Distribution of SB mapping reads across four distance intervals from the Sox2_-116 kb launch site (<10 kb, 10–100 kb, 100– 500 kb and 500 kb–2 Mb) for transposons carrying neutral_2 or Sox2 promoter cargo. Dots represent biological replicates. **(D)** Distribution of SB mapping reads across the same distance intervals as in (C) from the Nanog_1 launch site for transposons carrying neutral_1, Nanog promoter or synthetic promoter cargos. Dots represent different reporter constructs.

For the same neutral cargo, the fraction of all reintegrations that were local varied between launch sites (**Fig S7A**). This indicates that the launch site position contributes to the balance between local and distal reintegration. We therefore focused on the two launch sites for which multiple cargos were available: Sox2_-116kb and Nanog_1. At both launch sites, promoter and synthetic promoter cargos (see **Fig 3C**) generally showed a higher fraction of reintegrations *in trans* (**Fig 5B**) and a lower fraction of local reintegrations (**Fig S7B**) than the short neutral cargo. Thus, promoter-containing cargos were less likely to reintegrate on the same chromosome as the launch site.

In contrast, CBS-containing cargos were more strongly confined to the local region and showed fewer *trans* reintegrations than the neutral cargo at launch site Sox2_-116kb (**Fig 5B, Fig S7B**). This effect was observed both for CBS-only cargos and for cargos in which CBSs were combined with the *Sox2* promoter. Thus, cargo can shift the balance between local and interchromosomal SB transposition in either direction, depending its features.

We next asked whether cargo sequence also affects the location of reintegrations within the local 2 Mb region. Therefore, we subdivided local reintegrations into four distance classes: <10 kb, 10–100 kb, 100–500 kb, 500 kb–2 Mb. Because the 3D chromatin folding of the two launch sites causes a different local pattern of reintegration locations for the neutral cargo (see also **Fig 4H**), we compare between the cargos per launch site. Promoter-containing cargos were more enriched very close to the launch site than the neutral cargo. At launch site Sox2_-116kb, 16% of local reintegrations of the promoter cargo occurred within 10 kb of the launch site, compared to 6% for a short neutral cargo (**Fig 5C**). A similar pattern was observed at Nanog_1: cargos containing either a *Nanog* promoter or a synthetic promoter showed a higher fraction of very proximal reintegrations than the neutral cargo (**Fig 5D**). Within the synthetic promoter set, the two most active reporters showed more proximal reintegrations, suggesting the difference may be related to cargo activity, although this trend was not significant across all reporters (**Fig S7C**).

Together, these results show that SB cargo influences reintegration location at multiple scales. Promoter-containing cargos were less likely to remain within 2 Mb of the launch site, but among the reintegrations that remained local, they were enriched very proximal (<10 kb) to the launch site. In contrast, addition of CBS to the reporter cargo reduces the fraction of transpositions to other chromosomes. These findings indicate that, in addition to 3D contact, cargo sequence contributes to the genomic distribution of SB reintegrations.

## Discussion

Since its reconstruction from inactive fish transposon sequences (*1*), SB has been widely used for genome engineering, mutagenesis and gene delivery (*11*). Local SB transposition from a chromosomal launch site has been particularly powerful for the detailed study of individual genomic loci, either by mutagenesis or the insertion or reporter genes (*6, 8–13*). Here, we used defined chromosomal launch sites to dissect the determinants of genome-to-genome SB transposition in mouse embryonic stem cells. First, mobilization efficiency depends on both launch site context and cargo properties. Transcriptionally active launch sites and SB with larger or more transcriptionally active cargos have reduced transposition efficiency. Second, after mobilization, reintegration patterns are primarily shaped by spatial genome organization. For neutral cargos (i.e. without transcriptional or regulatory activity), local reintegration frequency closely follows three-dimensional contact with the launch site and shows little additional dependence on chromatin context at either the launch or reintegration site. However, other cargos can modify this local reintegration pattern and the frequency of transposition to other chromosomes. Together, these results show that the factors affecting how efficiently SB mobilizes are not the same as those determining where it reintegrates.

In the clonal assays, only 40–52% of excision-positive clones showed a detectable reintegration event, leaving a substantial fraction of excised transposons without a mapped insertion site. A similar phenomenon was reported previously (*3*). One possible explanation is transposition into the sister chromatid after DNA replication. In that case, one daughter cell will inherit a transposed copy of SB and one that may have remained in the launch location (thus counted as ‘no mobilization’), whereas the other daughter inherits none (counted as ‘no reintegration’). Auto-integration of the SB into its own cargo sequence may provide an additional explanation for lost transposons, although a previous study showed that increasing SB transposon size reduces productive transposition primarily through reduced excision, with auto-integration having a more modest contribution (*27*). Finally, SB transposons that fail to reintegrate before the next mitosis might end up in the cytoplasm and be lost.

In the nested-transposon system, we found that local SB transposition was positively associated with heterochromatic features such as LaminB1 and H3K9me3 at the launch site, while negatively associated with transcriptionally active and accessible chromatin. Our results are consistent with previous studies showing enhanced SB transposition after targeted heterochromatin formation by KRAB recruitment (*28*) or DNA methylation (*29, 30*). This might be explained by the chromatin compaction level, since DNA compaction was shown to increase transposition (*30*), presumably by facilitating the productive pairing of the two transposon ends by the transposase which is a necessary step for excision. Alternatively, there might be a preferential localization of SB transposase to heterochromatin (*28*).

The negative association between transcriptionally active chromatin and SB transposition, partially reflects the *absence* of heterochromatin. However, the negative linear relationship between transposition and euchromatic features like chromatin accessibility (measured by ATAC-seq), even at higher signal levels, suggests a direct effect of these features on transposition. We speculate that the binding of transcription factors or RNA polymerase II might interfere with mobilization. A direct effect of polymerase is consistent with the reduced transposition at transcribed loci not proximal to cis-regulatory elements. It is also supported by the reduced transposition of SB with an actively transcribed cargo, which directly causes an increase in local transcription factor and polymerase binding. Indeed, earlier work suggested that continuous binding of a DNA-binding protein inside the SB cargo reduces SB transposition (*28*). However, to our knowledge this is the first study that indicates an inhibitory effect of active chromatin features and transcription on transposition.

Experiments from single-launch-site cell lines revealed variability between launch sites in the fraction of reintegrations on other chromosomes. This could affect the number of local reintegrations assigned to each launch site in the nested-transposon experiments. However, the nested-transposon results agreed with observations from individual SB cell lines, where transcriptionally active cargo was associated with altered transposition efficiency. Although correlative, these findings support a model in which transposon occupancy and chromatin environment influence productive SB transposition.

Beyond mobilization efficiency, we found that reintegration pattern of SB with a neutral cargo closely followed three-dimensional contact with the launch site, which indicates that spatial proximity is a major determinant of the SB reintegration site. This is in agreement with qualitative observations from individual launch sites (*18, 31*), but the relationship was not previously quantified. The chromatin features of the excised transposon, including histone modifications or association to the nuclear lamina (LaminB1), do not affect these reintegration patterns, suggesting that they do not affect the mobility or reintegration ability of the excised transposon. In addition, the chromatin features of the reintegration sites do not affect SB reintegration frequency, since the reintegration pattern can be fully explained by contact with the launch site. A previous study found that SB reintegration sites are depleted in repressed chromatin (*31*), but the authors did not correct for the contact frequency with the SB launch site, so it is likely this depletion was a result of reduced contact between the heterochromatin region and the launch site. In addition, earlier studies using SB mobilized from a donor plasmid showed nearly random distribution across the genome without a preference for promoter-proximal or accessible regions (*32, 33*). Overall, these findings indicate that contact frequency alone can predict the distribution of SB reintegrations after excision, without effects of chromatin.

Since we previously observed an effect of CBSs on the direction of SB transposition, presumably by altering the genome folding before transposition (*19*), we investigated whether other cargo might also influence reintegration patterns. We found that promoter containing cargo increases the fraction of local reintegrations that are very proximal (<10 kb) to the launch site. It is probable that this is also due to the cargo changing 3D genome folding, since promoters (*34*) and RNA polymerase II (*35, 36*) are implicated in determining genome folding, and transcribed genes tend to form self-interacting domains (*37*). Paradoxically, besides the increase in very proximal reintegrations, promoter cargo also increases the fraction of transpositions to other chromosomes. It is unlikely that the reintegration of a single promoter into a euchromatic region affects genome folding on the chromosome scale. Instead, we speculate the promoter cargo might decrease SB reintegration efficiency, causing the transposon to search longer for an reintegration site and thus traveling further. Alternatively, the promoter-containing transposon might be attracted or trapped into transcriptionally active condensates that allows it to access other chromosomes more effectively. In contrast, CBSs in the cargos reduce the transposition to other chromosomes. It is possible that binding of CTCF and cohesin to the cargo reduces SB mobility, or even tethers it to the launch chromosome. Further research is needed to fully understand how cargo influences SB reintegration location.

The effects of launch site chromatin and cargo on SB transposition described here, can be used to inform experimental designs using local SB transposition, such as mutagenesis approaches. First, one could prioritize launch sites in heterochromatin or away from active *cis*-regulatory elements to improve transposition efficiency. Moreover, prediction of the distribution of reintegrations based on a 3D contact map can be used to select a launch site with sufficient reintegrations in a region of interest. Finally, it is important to consider the effect of the desired cargo. Taking the Sox2_-116kb launch site as an example, SB with a neutral cargo will result in around 23% of transposons relocated more than 10kb but less than 2Mb away from the launch site (40% transposed fraction * 60% local * 95% outside 10kb), while the Sox2P reporter cargo reduces this to 2.1% (5% transposed * 50% local * 85% outside 10kb). In general, to obtain a similar number of useable reintegrations, experiments with larger and more active cargo need to be scaled up.

Here we combined a large targeted nested-transposon experiment with reanalysis of SB transposition data generated for various projects to derive rules of SB transposition. The comparison of pre-existing data allowed us to compare many different conditions for these relatively large and labour-intensive experiments. However, this also means that perfectly matching conditions were missing in some comparisons. Future study designs could incorporate a set of cargos with matched transcriptional activity while varying in genomic size, to directly address the effect of both variables on transposition efficiency. In addition, future work could screen for cell perturbations that increase transposition frequency, to improve applications that require a high transposition efficiency. Finally, more functional studies could elucidate how the effects of chromatin context and cargo on transposition are mediated, or how cargo can affect transposition to other chromosomes.

## Acknowledgments

We thank the members of our laboratory for helpful comments and the NKI Genomics, Flow Cytometry, Animal Modelling Facility, and Research High-Performance Computing core facilities for excellent technical support. We thank Max Trauernicht, Chaitanya Rastogi and Harmen J. Bussemaker for designing and sharing the TP53 synthetic promoter sequence.

## Author Contribution

C.J.I.M., M.E. and B.v.S. conceptualized the study. M.E., C.J.I.M., L.D., M.V., J.O.Y.C. and L.E. performed wet-lab experiments. C.J.I.M. performed data analysis and developed code, with contributions from M.E., A.A.K., and V.H.F.S. M.E., C.J.I.M. and B.v.S. wrote the manuscript. B.v.S. supervised the project. B.v.S. and C.J.I.M. acquired funding.

## Data and materials availability

Laboratory notebooks and supplementary data (including plasmid sequences, FACS data, etc.) are available on Zenodo (10.5281/zenodo.23729914). Raw sequencing data is available on GEO under accession number GSE352692.

Code and supplementary files used as input for the scripts are available on Zenodo (10.5281/zenodo.23764413 and 10.5281/zenodo.23803168) and GitHub. All plasmids used will be made available upon request.

## Materials & Methods

### Experimental

#### Cell culture

Datasets generated from previously published F121/9 (CAST/EiJ × S129/Sv) female mouse embryonic stem cell (mESCs; RRID:CVCL_VC42) (*38*) experiments have been either cultured in Serum+Lif+2i, as previously described (*13*), or in serum free 2i+Lif media (*18*). WT78, WT79 and WT83 mESCs clones were initially cultured on feeder cells (irradiated mouse embryonic fibroblast (MEFs)) in Glasgow minimal essential medium (GMEM; Gibco, 21710-025) supplemented with 10% fetal calf serum (Thermo Fisher Scientific, 10270-106), 1% sodium pyruvate (Thermo Fisher Scientific, 11360-70), 1% MEM non-essential amino acids (Thermo Fisher Scientific, 11140-035), 1% penicillin–streptomycin (Thermo Fisher Scientific, 15070063), 100 μM β-mercaptoethanol, and 10 x 4 U leukemia inhibitory factor (LIF; esg1107, Millipore). MEFs were cultured in Glasgow minimal essential medium (GMEM; Gibco, 21710-025) supplemented with 10% fetal calf serum (Thermo Fisher Scientific, 10270-106), 1% sodium pyruvate (Thermo Fisher Scientific, 11360-70), 1% MEM non-essential amino acids (Thermo Fisher Scientific, 11140-035), and 1% penicillin–streptomycin (Thermo Fisher Scientific, 15070063). MEFs were irradiated with 20 Gy for 20 min and frozen in aliquots. Before using clones WT78, WT79 and WT83 for transposition experiments, they have been adapted to 2i+Lif media. The Sox2P_CDS reporter in Sox2_-161kb in *Sox2*::mCherry deleted cells were cultured in Serum+Lif+2i, as previously described (*13*). All other mESCs have been cultured in 2i+Lif as described previously (*18*). Cells were grown at 37 °C in a humidified incubator with 5% CO₂.

#### Plasmids

To generate pJOYC037, a short neutral Sleeping Beauty construct nested within PiggyBac transposon arms, the region spanning the left and right Sleeping Beauty transposon arms and an intervening loxP site was amplified by PCR from pJOYC032. Primers were designed to introduce an EcoRI restriction site at the 5′ end and a SpeI restriction site at the 3′ end of the amplicon (oJOYC-207 & oJOYC-208). The PCR product was digested with EcoRI and SpeI and cloned into the corresponding sites of the PiggyBac recipient plasmid pJOYC030, placing the Sleeping Beauty construct between PiggyBac transposon arms. The resulting plasmid, pJOYC037, was verified by Sanger sequencing.

The short (S, 557 bp) and long (L, 1593 bp) versions of the Nanog promoter were amplified from the genome using primers CMP011 or CMP012 with CMP013. These fragments were cloned into a recipient plasmid containing GFP, followed by a fragment of the human LBR gene (also used in (*39*), amplified with CMP008+CMP009) and the short sNRP-1 poly-A signal (also used in (*21*), amplified with CMP008 + CMP009) using Gibson assembly. F3 and FRT heterotypic Flp-recombinase sites were added by PCR (CMP016+CMP017) resulting in pCM002 (NanogP_S) and pCM003 (NanogP_L) respectively. To generate pCM008, the LBR fragment and sNRP-1 sNRP-1 poly-A signal in pCM002 were replaced by an SV40 poly-A signal (amplified from pME034, CMP044+MEP117) by PCR of pCM002 (CMP042 + CMP048) followed by Gibson assembly. pCM010 was generated by removing the LBR fragment from pCM002 by PCR (CMP042+CMP055) and self-ligation by Gibson assembly. The results from inserts pCM002, pCM008 and pCM010 are grouped as NanogP_S.

The synthetic promoter reporters contain the FRT/F3 sites in the opposite orientation from the Sox2 and Nanog promoter reporters used here. First intermediate plasmid pLE001 was generated from pCM010 by amplifying the Nanog promoter and GFP with a flanking F3 site (CMP143+CMP144) and the SV40 poly-A signal with flanking FRT (CMP145+CMP146), followed by Gibson assembly and blunt-cloning into a pZero vector. The synthetic promoters were ordered as gBlocks (Twist Biosciences) and cloned into an intermediate plasmid upstream of GFP. See **Table S4** for the reporter sequences, the TP53 reporter uses low affinity binding sites and is based on (*40*), the other synthetic promoters are from (*25*). The Nanog promoter and GFP in pLE001 were replaced by each of the synthetic promoters with GFP, by restriction digestion (AgeI and NdeI) and ligation, resulting in pLE005, pLE006, pLE007, pLE009 and pLE011.

To generate pMV001, a 2.5kb fragment of the superenhancer 5kb upstream of Nanog was PCR amplified from the genome (MVP001+MVP002), followed by a PCR to add FRT/F3 sites (MVP003+MVP004) and blunt cloned into the pZero vector. All gRNA and primer sequences can be found in **Table S5**.

#### Recombination-mediated cassette exchange (RMCE)

As described previously (*13, 18*), launch site cell lines were generated by first generating a founder cell line containing a HyTK dual-selection marker flanked by heterotypic FRT/F3 recombination sites and Sleeping Beauty inverted terminal repeats. Then, RMCE was used to replace the HyTK selection cassette with defined cargo constructs. The Nanog promoter reporters were inserted into the Nanog_1 HyTK cell line previously published (*18*), by transfecting 3 × 10^5 cells were transfected with 1.5 µg of the donor plasmid (pCM002, pCM003, pCM008 or pCM010) and 0.5 µg of a Flp recombinase expression plasmid (Addgene, 13787). Two days after transfection, ganciclovir was added to the medium at 2.5 µg/mL to select for HyTK-negative cells. Colonies were picked seven days after the start of selection and screened by PCR and Sanger sequencing.

The synthetic promoters (pLE005, pLE006, pLE007, pLE009 and pLE011) and the Nanog enhancer (pMV001) were inserted similarly, but into a genomically edited version of the Nanog_1 launch site cell line. For reasons unrelated to this manuscript, a destabilized Cas9 (*41*) driven by a constitutive CAG promoter was integrated in the Rosa26 locus in this launch site cell line before RMCE.

The cell lines with the neutral-1 insert (pLD037) in the Tdgf1_1, Nanog_1 and Fzd5_1 launch sites were previously published (*18*), as were all cell lines with inserts in the Sox2_-161kb and Sox2_-116kb launch sites (*13, 19*).

#### Establishment of cell lines from Sleeping Beauty hopping

Cell lines carrying reintegrated Sleeping Beauty cargo constructs were established by clonal isolation after Sleeping Beauty hopping from previously generated launch site cell lines. Tdgf1_3, Tdgf1_48 and Tdgf1_137 were isolated after hopping from the Tdgf1_1 launch site carrying pLD037 as cargo. Fzd5_108 was isolated after hopping from the Fzd5_1 launch site carrying pLD037. Nanog_31 and Nanog_37 were isolated after hopping from the Nanog_1 launch site carrying pLD037. Sox2-inSCR was isolated after hopping from the Sox2-116 kb launch site carrying the Sox2P reporter cargo. All launch site coordinates are listed in **Table S1.**

#### Establishment of clones carrying multiple launch sites

F121/9 female mouse embryonic stem cells (mESCs; CAST/EiJ × S129/Sv; RRID:CVCL_VC42) (*38*) were used to establish clones carrying multiple Sleeping Beauty launch sites. Cells were co-transfected with a PiggyBac expression vector carrying an NGFR selection marker (pJOYC033_hPGK_PB_IRES_NGFR) and a short neutral Sleeping Beauty launch site construct (pJOYC037_HE227_PB_SB) at a 1:3 plasmid ratio. For each transfection, 2 × 10^6 cells were lipofected with 4 µg total plasmid DNA and 12 µl Lipofectamine 2000 (Invitrogen, 11668019).

Two days after transfection, NGFR-positive cells were enriched by magnetic-activated cell sorting using MACS MS columns (130-042-201, Miltenyi Biotec) and LNGFR MicroBeads (130-091-330, Miltenyi Biotec), according to the manufacturer’s instructions. During transfection and for two weeks after transfection, cells were cultured in 60% BRL-conditioned medium supplemented with β-mercaptoethanol and LIF. Cells were then transferred to mouse embryonic fibroblast feeder cultures.

Single-cell clones were isolated by limiting dilution. Three clones, designated WT78, WT79 and WT83, were selected for further analysis. Cells were adapted from feeder culture to 2i medium for two weeks. Afterwards, launch sites (i.e. PB integrations) were mapped by Tn5-based mapping (see ‘Tagmentation mapping’).

#### Sleeping Beauty transposition

Besides the re-analysis of previously published data, Sleeping Beauty transposition was induced for this study in several cell lines. Sleeping Beauty transposition was performed as described previously, with minor modifications. Briefly, 1 to 2.5 million cells were transfected using, per 1 × 10^6 cells, 5 µg of the bicistronic plasmid pME07, encoding SB100X transposase and human nerve growth factor receptor (NGFR), using 15 µl Lipofectamine 2000 (Invitrogen, 11668019). As a control, 5 × 10^5 cells were transfected with 2.5 µg of the analogous bicistronic plasmid pLD042, encoding PiggyBac transposase and LNGFR. Thirty hours after transfection, NGFR-positive cells were enriched by magnetic-activated cell sorting using MACS MS columns (130-042-201, Miltenyi Biotec) and NGFR MicroBeads (130-091-330, Miltenyi Biotec), according to the manufacturer’s instructions.

For the Nanog_31, Nanog_37, Tdgf1_3, Tdgf1_137 and Fzd5_108 launch sites carrying the short neutral insert neutral_1 (pLD037), pools were harvested for tagmentation mapping at multiple timepoints between 4 and 11 days after transfection. Since these are subpools from the same transfection, they are combined in the data analysis. For the clones carrying multiple SB launch sites (WT78, WT79, WT83), and for the pMV001, pCM002, pCM003, neutral_1 insert in the Nanog_1 launch site the entire pool was harvested either 7 or 10 days after transfection. For the Sox2P reporter in the Sox2_inSCR launch site 0.2 million cells were set apart 7 days after transfection and harvested after 4 more days of expansion. The same protocol was followed for the Sox2P_CDS reporter in Sox2_-161kb in *Sox2*::mCherry deleted cells (see (*14*) for generation of the cell line).

The cell lines containing *Nanog* reporters pCM008 and pCM010 and the synthetic promoters (pLE005, pLE006, pLE007, pLE009 and pLE011) were also CRISPR-edited to introduce random barcodes inside the GFP sequence. These barcodes are not used in this study, but the procedure will be briefly described. For the pCM008 and pCM010 inserts in the Nanog_1 launch site, cells were first co-transfected with a plasmid containing Cas9 and a gRNA targeting GFP (**Table S5**) and a single stranded repair template which will introduce a 12 nucleotide random sequence (oCM005). Cells were treated for 5 days with 1 µM DNA-PK inhibitor M3814 (M3814, MCE, HY-101570). Pools of GFP-negative cells were sorted to enrich for barcode insertion and then SB transposition was induced in both sorted and unsorted cells. Three to eight days after transposase transfection, three pools of 500 cells were generated by serial dilution. Since these subpools are derived from the same transfection, they are combined in the data analysis. For the cell lines with synthetic promoters, barcode integration was induced after transposition. Eight days after transposase transfection, the cell pools were transfected with a plasmid containing the gRNA targeting GFP (but no Cas9) and a single stranded repair template which will introduce a 16 nucleotide random sequence (oCM012). Cells were treated for 4 days with 1 µM DNA-PK inhibitor M3814, and with 0.5 µM Shield-1 (AOB1848) to stabilize the constitutively expressed Cas9. Cell pools were harvested around 10 days after the last transfection.

#### Assessment of excision efficiency

To assess Sleeping Beauty excision efficiency, single cells were sorted by flow cytometry into 96-well plates one week after initiation of Sleeping Beauty hopping. Crude genomic DNA extracts were prepared from each clone using 25 µl DirectPCR lysis reagent (302-C, Viagen Biotech) supplemented with 100 µg/mL proteinase K. Lysates were incubated at 55 °C for 150 min, followed by heat inactivation at 85 °C for 45 min.

Sorted clones were screened by PCR using primers flanking the launch site, with 2–3 µl crude lysate used as template. Successful excision of the transposon resulted in amplification of a shorter PCR product compared with the non-excised launch site allele. Excision efficiency was calculated as the fraction of clones with detectable transposon excision among all analyzed clones. The number of analyzed and excised clones for each launch site cell line is listed in **Table S2**.

#### Assessment of reintegration efficiency

To assess detectable reintegration efficiency, selected clones with confirmed transposon excision were subjected to Tn5-based mapping (see ‘Tagmentation mapping’). Not all clones with detectable excision were processed for mapping. Clones with one or more mapped Sleeping Beauty reintegrations were classified as “excised and reintegrated”, whereas clones where the tagmentation mapping resulted in no mapped reintegrations, were classified as “excised, no detectable reintegration”.

Detectable reintegration efficiency was calculated as the fraction of mapped excised clones with at least one detected reintegration event. The number of excised clones subjected to Tn5-based tagmentation mapping and the number of clones with detected reintegrations are listed in **Table S2.**

#### Flow cytometry

Reporter fluorescence was measured by flow cytometry on a BD LSRFortessa. GFP expression from synthetic promoters, NanogP_S and NanogP_L cargos inserted at the Nanog_1 launch site was measured using the eGFP channel with 488 nm excitation and a 530/30 emission filter. Single cells were gated in FlowJo, and fluorescence data were exported as .fcs files for downstream analysis in R using the flowCore package (*42*).

Reporter measurements of the Sox2P cargo at the Sox2-116 kb launch site in CBS-WT, ΔCBS-in-SCR and ΔCBS-DS-SCR genomic backgrounds were described previously (*19*). Reporter measurements of Sox2P and Sox2P_CDS cargos at the Sox2_-161 kb launch site, with or without deletion of endogenous *Sox2*, were described previously (*13*).

#### Tagmentation mapping

All Sleeping Beauty reintegrations were mapped using the TagMap protocol (*20*), as described previously (*13*), with slight variations. Sleeping Beauty reintegration sites from the Fzd5_1, Nanog_1 and Tdgf1_1 launch site cell lines carrying pLD037 were mapped using the forward enrichment primer only. Reintegration sites from the Fzd5_108, Nanog_31, Nanog_37, Tdgf1_137, Tdgf1_3 and Tdgf1_48 cell lines carrying pLD037 were mapped using both forward and reverse enrichment primers, as described previously (*18*). For these samples, 3 µL of tagmented genomic DNA (corresponding to 12.5 ng) was used as input for linear amplification.

The neutral_1, pCM002 and pCM003 inserts in launch site Nanog_1 were mapped in two technical replicates, with either one or ten tagmentation reactions (each containing 100 ng genomic DNA), of which either 3 µL (12.5 ng) or 10 µL (41.7 ng) per reaction was used as input for linear amplification. For the synthetic promoters in launch site Nanog_1, the Sox2P reporter in the launch site Sox2_inSCR, and the Sox2P_CDS reporter in Sox2_-161kb in *Sox2*::mCherry deleted cells, ten 100 ng tagmentation reactions were performed, of which 10 µL (41.7 ng) per reaction was used as input for linear amplification. For the pMV001 insert in the Nanog_1 locus, ten 100 ng tagmentation reactions were performed, of which 3 µL (12.5 ng) per reaction was used as input for linear amplification. For the pCM008 and pCM010 inserts in launch site Nanog_1, one 100 ng tagmentation reactions was performed per pool (six total per insert), of which 7 µL (29.1 ng) per reaction was used as input for linear amplification using an alternative linear amplification primer (CMP070) which amplifies from inside the GFP gene. PiggyBac integration sites in clones carrying multiple launch sites were mapped using PiggyBac-specific primers instead of Sleeping Beauty-specific primers. TAC0099 and TAC0006 were used as reverse and forward enrichment primers, respectively. TAC0161 and TAC0110 were used as reverse and forward primers for PCR1, respectively. For these samples, 3 µL (12.5 ng) of tagmented genomic DNA was used as input for linear enrichment.

#### pA-DamID

pA-DamID on LaminB1 was performed on wildtype F121/9 (CAST/EiJ × S129/Sv) female mESCs grown in serum free 2i+Lif media. pA-DamID was performed as described previously (*43*),using 1:100 LaminB1 antibody (Abcam, ab16048, rabbit). The library preparation was performed as in (*44*). To process pA-DamID data, we used a previously published pipeline (*18*), aligning reads to the mouse reference genome (release M23, mm10) (*45*).

### Computational

#### Mapping integrations

We used the previously published pipeline (*18*) to map PB and SB integrations to a mouse reference genome (Release M23 – mm10). Because the Nanog_1 launch site contained additional sequences surrounding the SB cassette (*18*), Nanog_1 experiments were mapped to an *in silico* genome containing those surrounding sequences and integrations were lifted over to the reference genome. For published dataset, the processed mapping data as previously published was used. We only used mapping data from clones and cell pools that were not selected for any transposition-related phenotype, to obtain an unbiased representation of the Sleeping Beauty transposition behavior. **Table S6** outlines all datasets, generated here or previously published. The Sox2_-116kb cell line with neutral_2 insert was named the CRE6 −116 kb in (*13*). For the analysis of integration locations, biological replicates were merged.

#### Transposed fraction

To determine the transposed fraction from SB mapping in pools, we trimmed the SB arm and Tn5 adaptor sequences from all raw reads containing the expected SB sequence. Of all trimmed reads of at least 16 nt, the reads not matching the corresponding SB-flanking sequence in the launch site (allowing 2 mismatches) were assigned as transposed. Only samples with at least 15 untransposed reads were included. When mapping was performed for both SB arms on one sample, the mean of the two transposed fractions was used.

#### Relative local transposition

To determine the relative local transposition per launch site for cell lines with multiple launch sites, we first counted per sample the unique reintegrations within 2 Mb of each mapped launch site (i.e. PB integration). This number was normalized to the mean reintegration count of all launch sites per cell line per sample, giving the relative local transposition, and then averaged between the two biological replicates. For the comparison to chromatin marks, launch sites on sex chromosomes, in poorly mappable regions and those overlapping the ENCODE blacklist were removed (see ‘Quantification of reintegration events’).

#### Interchromosomal transposition

To determine the fraction of reintegrations on other chromosomes, we used a more stringent filtering of at least two mapping reads per reintegration. This is necessary because individual off-target mapping reads occur distributed across the genome, and therefore the fraction of reintegrations is sensitive to sequencing library quality when single mapping reads are counted as reintegrations. More stringent filtering for three or more mapping reads does not change the estimated fraction of trans hopping, but does drastically decrease the number of reintegrations that can be analysed (data not shown), therefore a threshold of at least two mapping reads was used. Samples with less than ten filtered reintegrations were excluded.

#### Quantification of reintegration events

In experiments where SB reintegrations were mapped in a highly complex pool of transposase transfected cells, a unique reintegration site can represent multiple independent reintegration events in that location. This causes a slight bias, where preferred reintegration sites (i.e. very close to the launch site) can be slightly underrepresented. In these experiments with highly complex pools, the mapping of the reintegration sites is relatively sparse: many reintegrations are only mapped by a single mapping read. Because there is no saturation in the unique mapping reads, the number of unique mapping reads reflects the abundance of the reintegration in the harvested cell pool and thus – on average – reflects the abundance of the reintegration event upon transposition. Therefore, we used the fraction of unique mapping reads as a proxy for the fraction of reintegration events when analysing the location of local Sleeping Beauty reintegrations. The data from inserts pCM008 and pCM010 were not included in the assessment of location reintegration locations, because those integrations were mapped after expanding 500 cell pools, and thus the read distribution may not reflect the original distribution of transposition sites anymore.

#### Motif reintegration sites

To determine the sequence motif at the SB reintegration sites, we used getSeq from the Biostrings package (*46*) to extract the 14 bp sequence surrounding the mapped reintegration coordinate. Next, we used the ggseqlogo package (*47*) to obtain the sequence motif. To compare the frequency of the flanking T and A (at +3 and −3) to their occurrence in the genome, we randomly sampled 10 million TA sites in the genome and classified them into the four possible motif versions (ANNTANNŢ BNNTANNT, ANNTANNV, BNNTANNV).

#### Quantification chromatin context

To characterize the chromatin context of the Sleeping Beauty launch sites, we collected 15 published mESC datasets (**Table S3**) and newly generated LaminB1 pA-DamID data. For the transient transcriptome sequencing (TT-seq) we used the original processed bigwig files, H3K9me3 pA-DamID we processed as the LaminB1 pA-DamID, for all other datasets we downloaded the uniformly reprocessed data from Cistrome v2.0 (*48*). Cistrome ChIP-seq tracks are not normalized to input, so to prevent biases from variation in genome mappability we excluded regions with unusually low or high mappability. Therefore we processed the raw fastq files from two deeply sequenced, single-end sequencing ChIP-seq input datasets (*49, 50*) according to the Cistrome processing pipeline (*48*) to obtain comparable bigwig tracks. To compare the feature signals to their genome-wide distribution, we used the bedtools function shuffle (*51*) to generate a set of one million random genomic coordinates (background set).

To quantify the signal around any coordinate (a launch site or a background set coordinate), we used a custom Snakemake pipeline available at https://github.com/vansteensellab/TRIP_local_context. In short, the average signal within a window (500 bp, 2 kb, 5 kb, 10 kb) around the launch site was calculated using DeepTools’s multiBigwigSummary (v3.5.2) (*52*). For TT-seq, the signal from forward and reverse mapping are added. Bedtools (v2.31.1) (*51*) was used to retrieve features near the launch sites, testing for overlap to genes of the GENCODE annotation (M21) (*53*) and distance to the closest DNase peak (ENCFF048DWN, (*54*)). For all features collected from cistrome, we added as a pseudocount the lowest value in the background set multiplied by 73 (this corresponds to half of one mapped read in the window). Afterwards, all features were log2-transformed (except the pA-DamID data, which is already in log-space).

After removing all coordinates on sex chromosomes, we determined the distribution of signal for the ‘input’ samples and removed all coordinates with input signal outside the range mean ± 2*SD (standard deviation) in either input sample, since these correspond to either poorly mappable regions or other mapping artifacts. Windows with any overlap to the ENCODE blacklist (*55*) were also excluded. The mean and standard deviation of feature signal across the remaining background set were used to calculate a z-score for all launch sites.

The chromatin context of reintegration windows is calculated in a similar way, centering the quantification on the middle of the 10 kb reintegration window instead of the launch site, and classifying z-score > 0 as high signal.

#### Quantification 3D contact

To correlate reintegrations to 3D contact with the launch site, we used the GENOVA package (*56*) to create virtual 4Cs from published microC data (*26*), at 10 kb or 25 kb resolution, using the launch sites as viewpoints. This analysis was performed on the same set of filtered launch sites obtained in the analysis of the chromatin context (seeQuantification chromatin context). Genomic bins without mapped reintegrations or with a contact value of 0 were removed, and the same filtered bins were used for the correlation between reintegrations and genomic distance. For the combined analysis of contact and chromatin state at the reintegration location, bins were also filtered for proper chromatin signal (as described above). In addition, launch sites with less than 30% high or low signal bins in the surrounding 2 Mb were excluded.

#### Flow cytometry

Flow cytometry data were analyzed in R using the flowCore package (*42*). For measurement of GFP-expressing reporters in the Nanog_1 launch site, the median fluorescence intensity was calculated for each sample. Autofluorescence was corrected by subtracting the median GFP fluorescence of a non-fluorescent wild-type control sample measured in the same experiment. For the synthetic promoters, autofluorescence-corrected median GFP values were then normalized to the random reporter and reported as relative GFP expression.

For Sox2P reporter measurements, the median fluorescence intensity was calculated for each fluorophore and each clone. Autofluorescence was corrected by subtracting the corresponding median fluorescence intensity measured in an mESC control cell line lacking fluorescent proteins. Corrected mTurquoise2 fluorescence, reflecting Sox2P reporter activity, was then normalized to the corrected eGFP fluorescence of the same sample to account for global differences in Sox2 expression (the endogenous *Sox2* gene on the other allele is tagged with eGFP). To correct for day-to-day variation in absolute fluorescence intensities, these normalized values were divided by the value measured for the original Sox2_-116 kb_Sox2P cell line on the same day. The resulting values were reported as relative Sox2P reporter expression.

## Supplementary material

### Supplementary tables

**Table S1.** Genomic coordinates of launch sites used in this study.

| Launch Region | Launch site name | Chromosome | Start coordinate |
|---|---|---|---|
| Fzd5_region | Fzd5_1 | chr1 | 64993495 |
| Fzd5_region | Fzd5_108 | chr1 | 79293540 |
| Tdgf1_region | Tdgf1_1 | chr9 | 110927616 |
| Tdgf1_region | Tdgf1_3 | chr9 | 109077492 |
| Tdgf1_region | Tdgf1_48 | chr9 | 109626920 |
| Tdgf1_region | Tdgf1_137 | chr9 | 110925102 |
| Nanog_region | Nanog_1 | chr6 | 122653234 |
| Nanog_region | Nanog_31 | chr6 | 143694588 |
| Nanog_region | Nanog_37 | chr6 | 109000999 |
| Sox2_region | Sox2_-161kb | chr3 | 34598479 |
| Sox2_region | Sox2_-116kb | chr3 | 34643960 |
| Sox2_region | Sox2_inSCR | chr3 | 34754848 |
| Sox2_region | Sox2_+715kb | chr3 | 35478413 |

**Table S2.** Number of clones screened for Sleeping Beauty excision and reintegration.

|  | Of all clones, based on PCR screen |  |  | Of excised clones, based on mapping |  |  | Of all clones, based on PCR screen and mapping combined |  |  |
|---|---|---|---|---|---|---|---|---|---|
| Launch pad | Analyzed clones | Excised clones | Fraction excised | Excised clones processed for mapping | Clones with detected reintegration | Fraction reintegrated | Fraction excised not reintegrated | Fraction excised and reintegrated | Fraction no-excision |
| Tdgf1_1 | 183 | 88 | 0.48 | 85 | 34 | 0.4 | 0.29 | 0.19 | 0.52 |
| Nanog_1 | 142 | 61 | 0.43 | 65 | 33 | 0.51 | 0.21 | 0.22 | 0.57 |
| Fzd5_1 | 164 | 65 | 0.40 | 64 | 30 | 0.47 | 0.21 | 0.19 | 0.60 |
| Tdgf1_48 | 290 | 104 | 0.36 | 52 | 30 | 0.58 | 0.15 | 0.21 | 0.64 |
| Tdgf1_3 | 313 | 48 | 0.15 | 23 | 12 | 0.52 | 0.07 | 0.08 | 0.85 |

**Table S3.** Datasets used for chromatin feature quantification.

| **Feature** | **Cistrome ID** | **GEO sample** | **GEO series** | **citation** |
|---|---|---|---|---|
| CTCF | 5933 | GSM747535 | GSE30203 | (*57*) |
| EZH2 | 9082 | GSM590133 | GSE23943 | (*58*) |
| H3K27ac | 66760 | GSM2416832 | GSE89209 | (*59*) |
| H3K27me3 | 74535 | GSM2416833 | GSE89209 | (*59*) |
| H3K36me3 | 9076 | GSM590120 | GSE23943 | (*58*) |
| H3K4me1 | 74534 | GSM2416834 | GSE89209 | (*59*) |
| H3K4me3 | 46553 | GSM769008 | GSE49847 | (*60*) |
| H3K9me3 |  | GSM7977244, GSM7977250, GSM7977256, GSM7977262 | GSE250306 | (*18*) |
| MED1 | 2857 | GSM560348 | GSE22557 | (*61*) |
| p300 | 74532 | GSM2433946 | GSE89209 | (*59*) |
| POLR2A | 46422 | GSM918749 | GSE49847 | (*60*) |
| RAD21 | 85637 | GSM2870307 | GSE74055 | (*62*) |
| TTseq |  | GSM6373843, GSM6373844 | GSE209526 | (*63*) |
| ATAC | 80533 | GSM2651155 | GSE99746 | (*64*) |
| DNase | 58283 | GSM1657364 | GSE67867 | (*65*) |
| input_1 |  | GSM4087828 | GSE137193 | (*50*) |
| input_2 |  | GSM3084076 | GSE112806 | (*49*) |
| LaminB1 |  |  |  | Generated in this study |

**Table S4.**
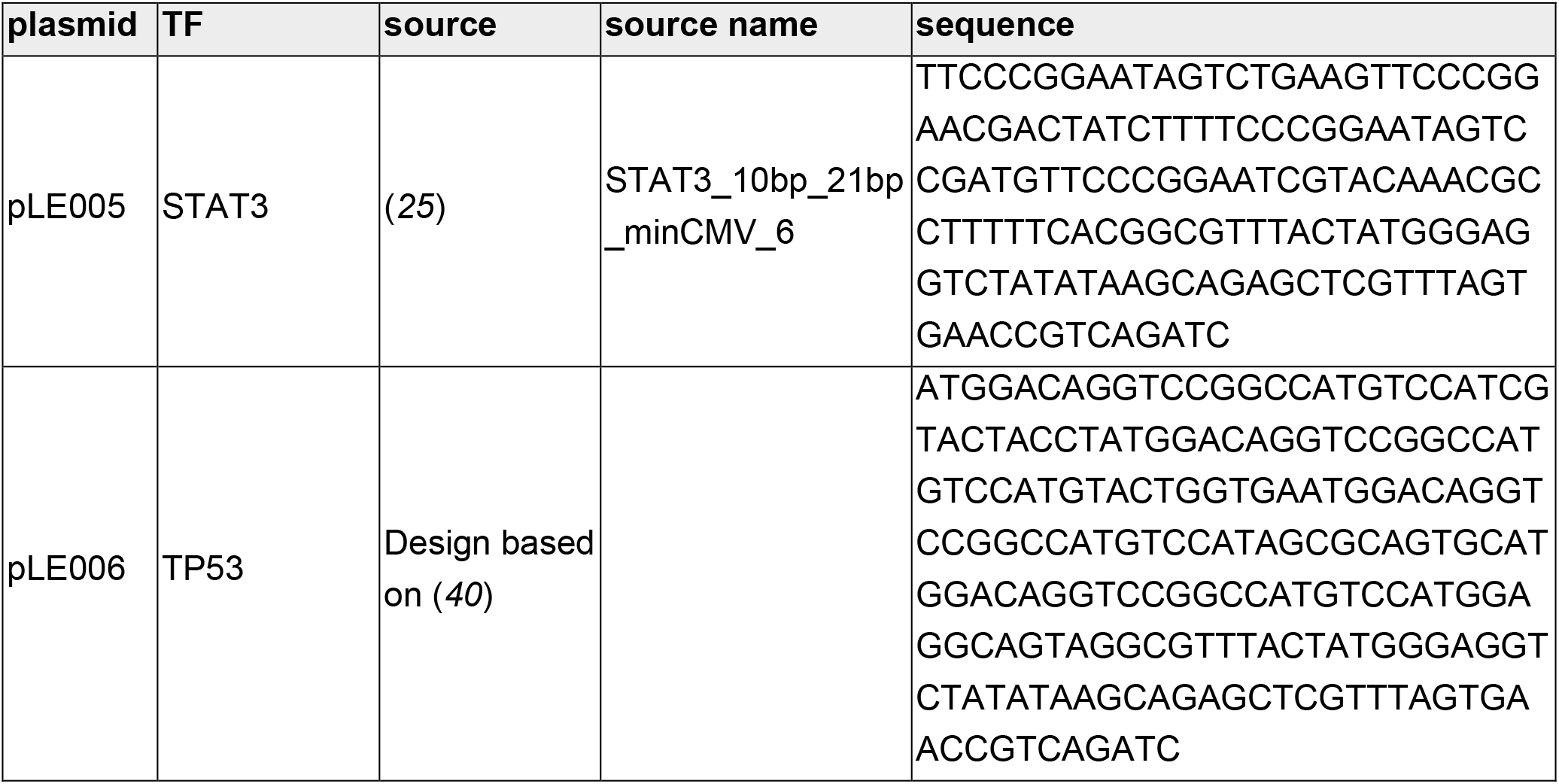
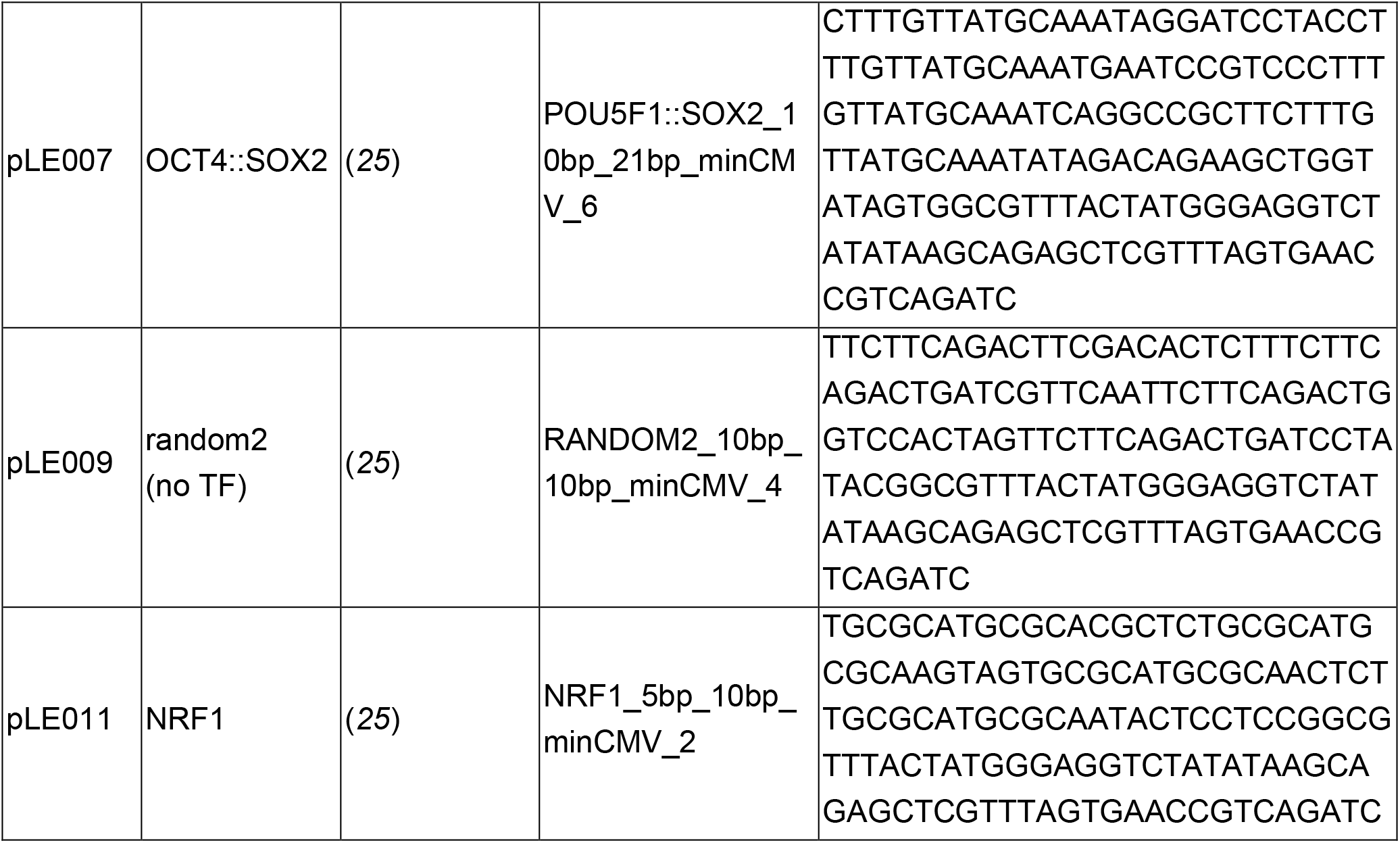
Sequence of synthetic promoter reporters.

**Table S5.**
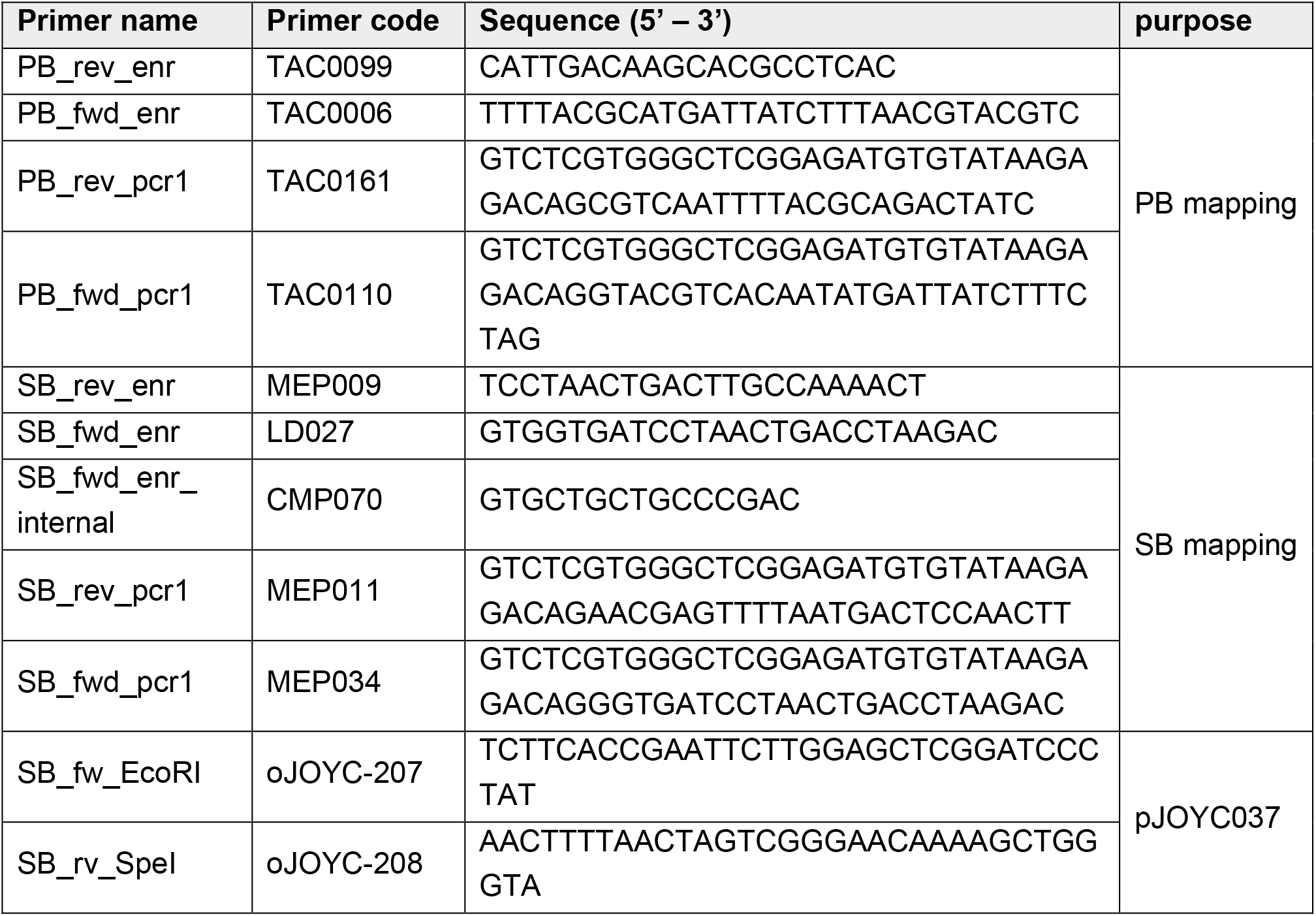
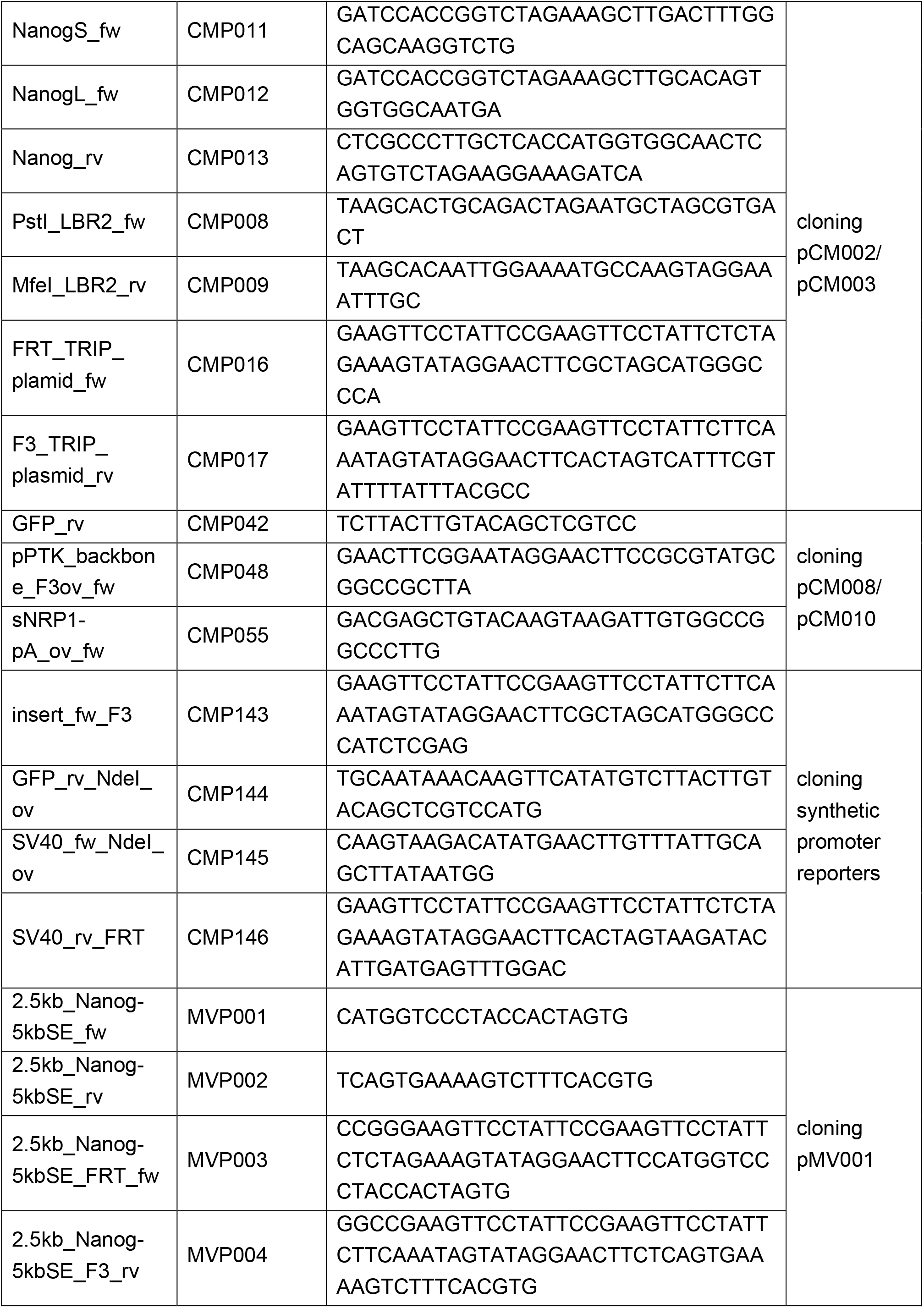
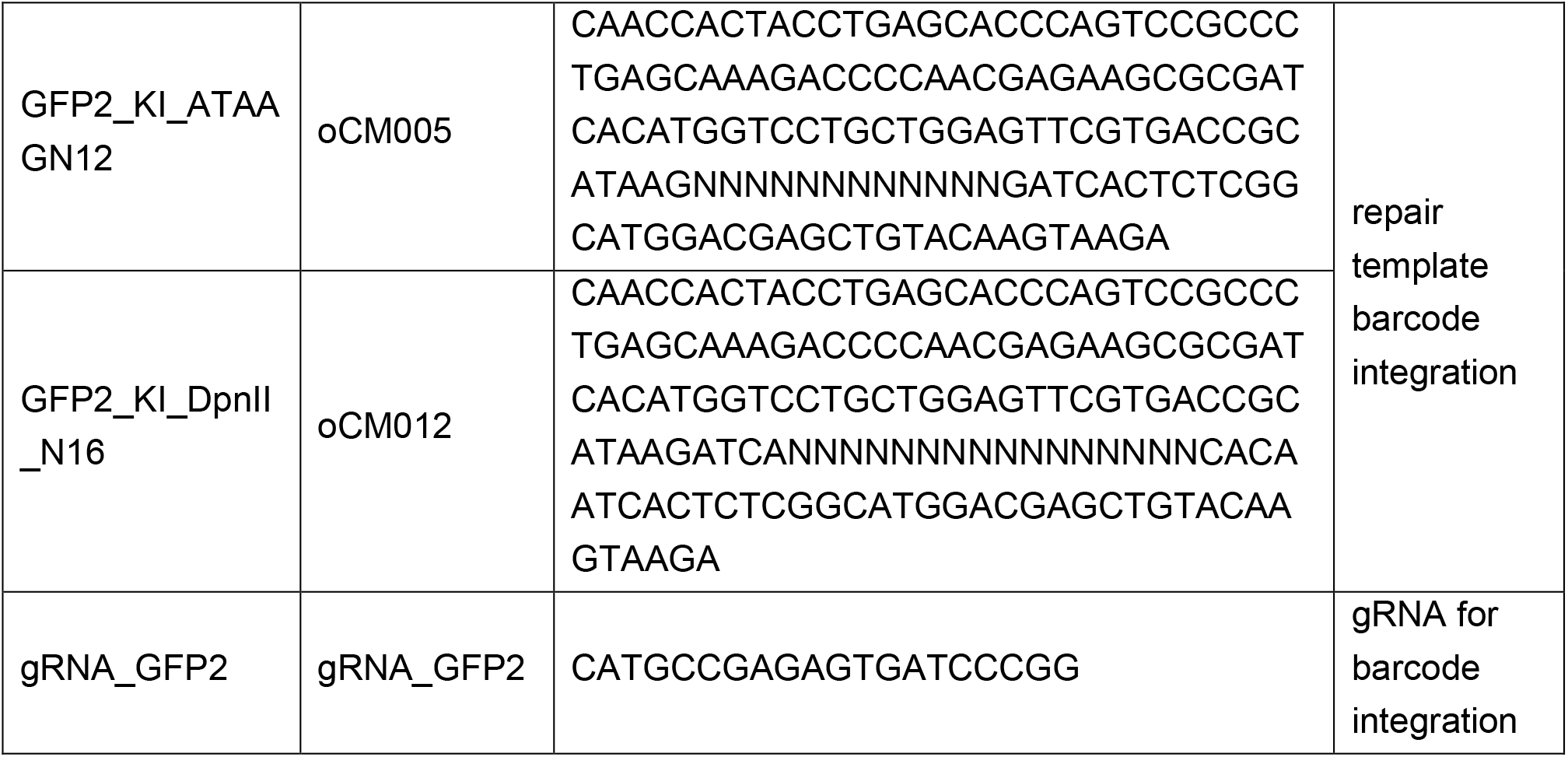
List of primers and gRNAs used in this study.

**Table S6.**
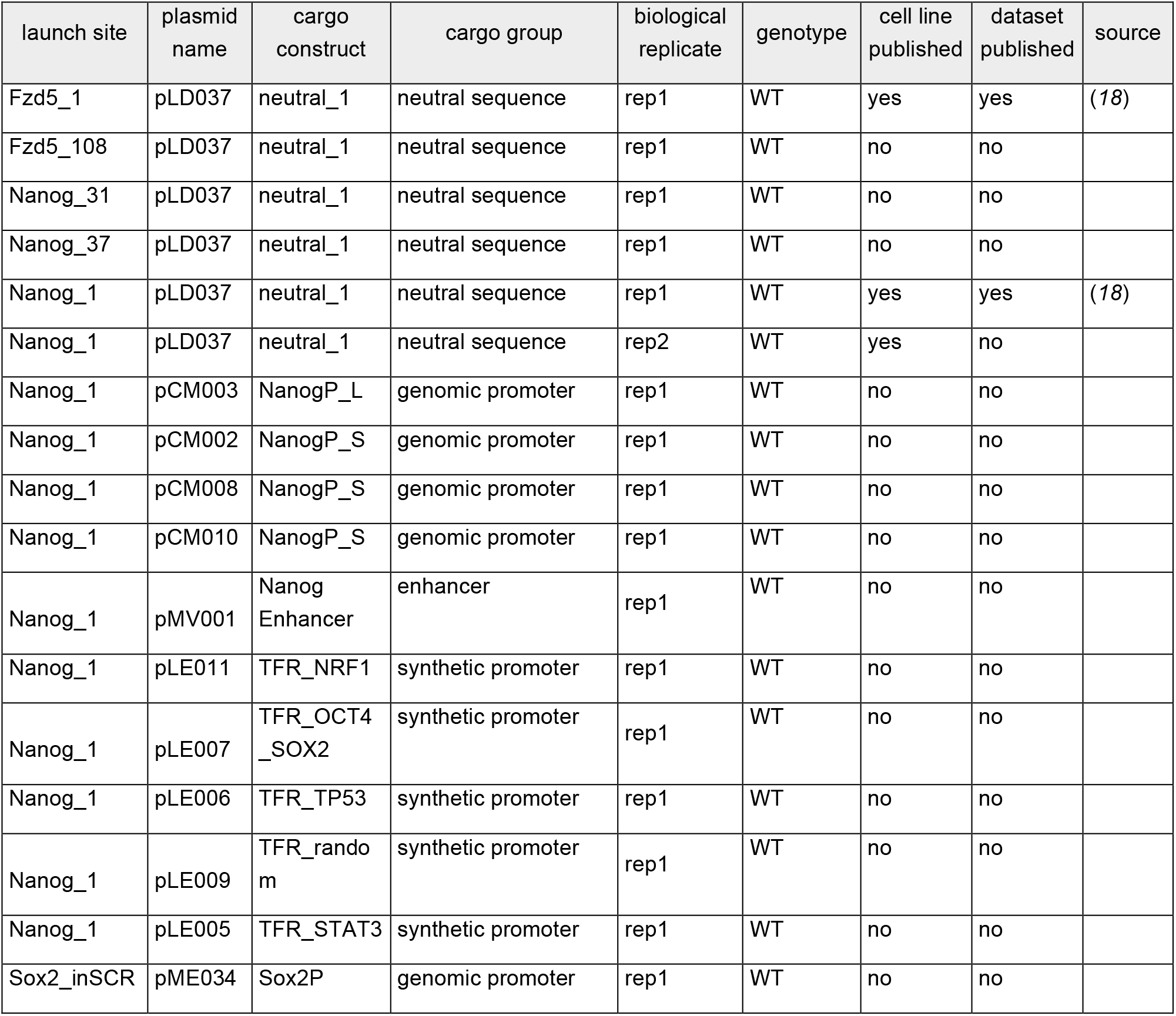
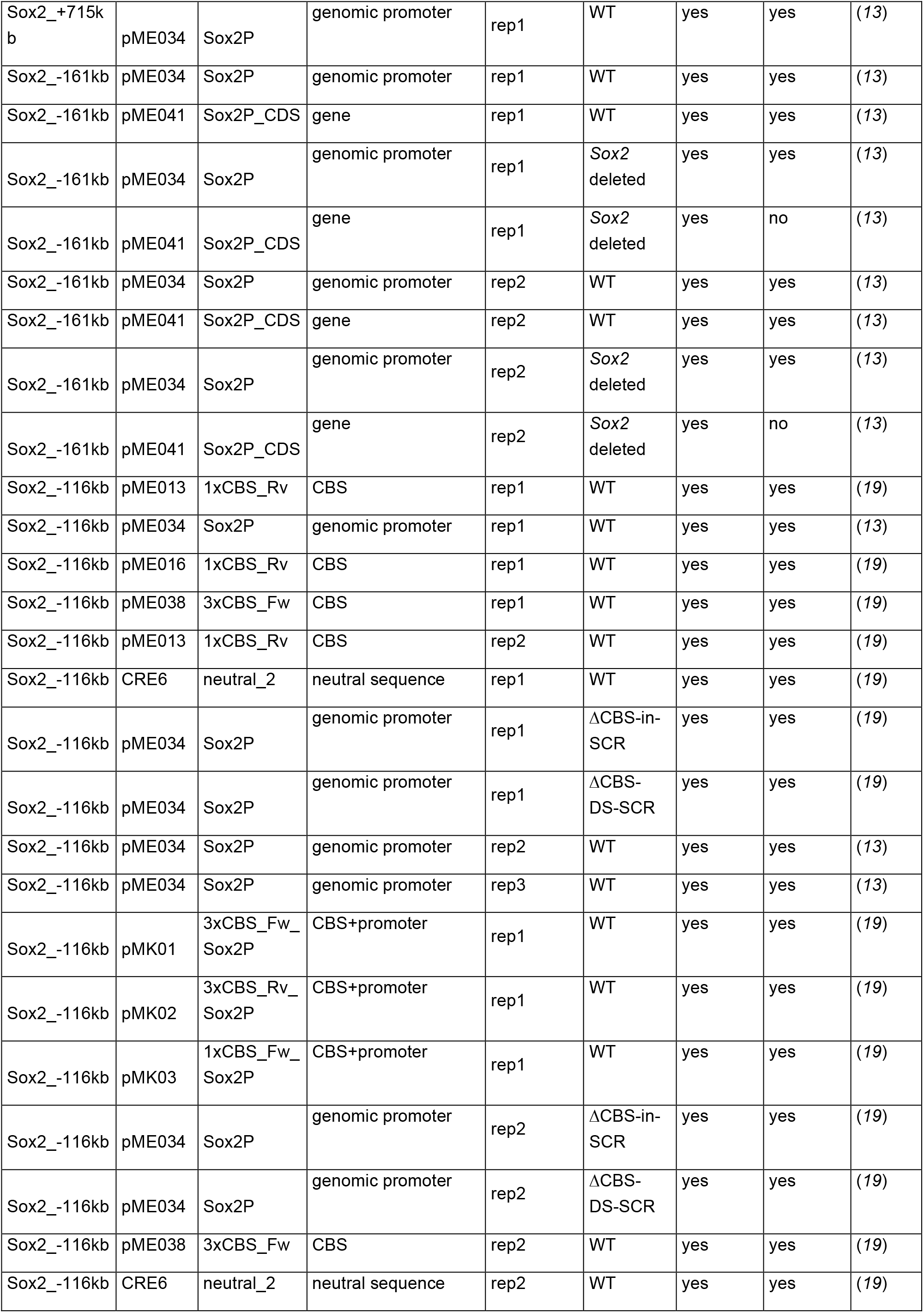
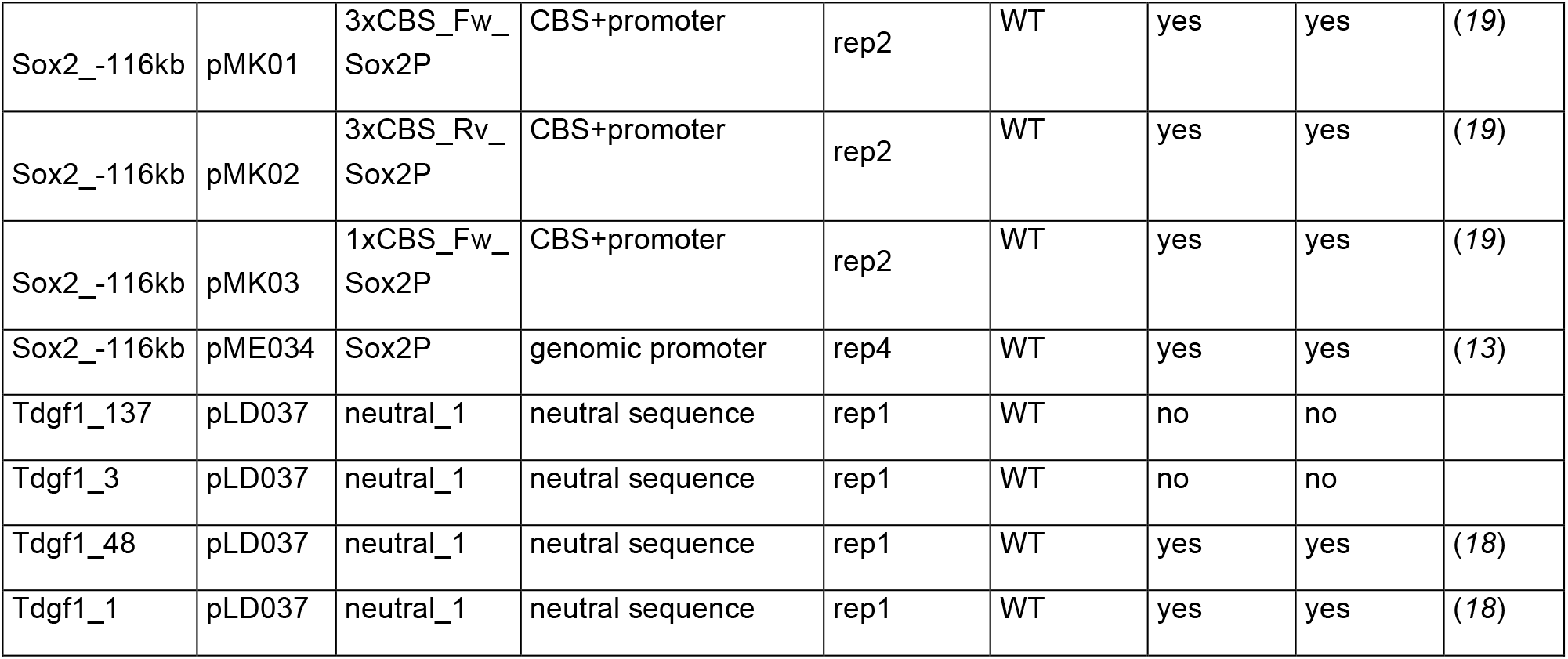
Overview of previously published and newly generated datasets.

| launch site | plasmid name | cargo construct | cargo group | biological replicate | genotype | cell line published | dataset published | source |
|---|---|---|---|---|---|---|---|---|
| Fzd5_1 | pLD037 | neutral_1 | neutral sequence | rep1 | WT | yes | yes | (*18*) |
| Fzd5_108 | pLD037 | neutral_1 | neutral sequence | rep1 | WT | no | no |  |
| Nanog_31 | pLD037 | neutral_1 | neutral sequence | rep1 | WT | no | no |  |
| Nanog_37 | pLD037 | neutral_1 | neutral sequence | rep1 | WT | no | no |  |
| Nanog_1 | pLD037 | neutral_1 | neutral sequence | rep1 | WT | yes | yes | (*18*) |
| Nanog_1 | pLD037 | neutral_1 | neutral sequence | rep2 | WT | yes | no |  |
| Nanog_1 | pCM003 | NanogP_L | genomic promoter | rep1 | WT | no | no |  |
| Nanog_1 | pCM002 | NanogP_S | genomic promoter | rep1 | WT | no | no |  |
| Nanog_1 | pCM008 | NanogP_S | genomic promoter | rep1 | WT | no | no |  |
| Nanog_1 | pCM010 | NanogP_S | genomic promoter | rep1 | WT | no | no |  |
| Nanog_1 | pMV001 | Nanog Enhancer | enhancer | rep1 | WT | no | no |  |
| Nanog_1 | pLE011 | TFR_NRF1 | synthetic promoter | rep1 | WT | no | no |  |
| Nanog_1 | pLE007 | TFR_OCT4_SOX2 | synthetic promoter | rep1 | WT | no | no |  |
| Nanog_1 | pLE006 | TFR_TP53 | synthetic promoter | rep1 | WT | no | no |  |
| Nanog_1 | pLE009 | TFR_random | synthetic promoter | rep1 | WT | no | no |  |
| Nanog_1 | pLE005 | TFR_STAT3 | synthetic promoter | rep1 | WT | no | no |  |
| Sox2_inSCR | pME034 | Sox2P | genomic promoter | rep1 | WT | no | no |  |
| Sox2_+715kb | pME034 | Sox2P | genomic promoter | rep1 | WT | yes | yes | (*13*) |
| Sox2_-161kb | pME034 | Sox2P | genomic promoter | rep1 | WT | yes | yes | (*13*) |
| Sox2_-161kb | pME041 | Sox2P_CDS | gene | rep1 | WT | yes | yes | (*13*) |
| Sox2_-161kb | pME034 | Sox2P | genomic promoter | rep1 | *Sox2* deleted | yes | yes | (*13*) |
| Sox2_-161kb | pME041 | Sox2P_CDS | gene | rep1 | *Sox2* deleted | yes | no | (*13*) |
| Sox2_-161kb | pME034 | Sox2P | genomic promoter | rep2 | WT | yes | yes | (*13*) |
| Sox2_-161kb | pME041 | Sox2P_CDS | gene | rep2 | WT | yes | yes | (*13*) |
| Sox2_-161kb | pME034 | Sox2P | genomic promoter | rep2 | *Sox2* deleted | yes | yes | (*13*) |
| Sox2_-161kb | pME041 | Sox2P_CDS | gene | rep2 | *Sox2* deleted | yes | no | (*13*) |
| Sox2_-116kb | pME013 | 1xCBS_Rv | CBS | rep1 | WT | yes | yes | (*19*) |
| Sox2_-116kb | pME034 | Sox2P | genomic promoter | rep1 | WT | yes | yes | (*13*) |
| Sox2_-116kb | pME016 | 1xCBS_Rv | CBS | rep1 | WT | yes | yes | (*19*) |
| Sox2_-116kb | pME038 | 3xCBS_Fw | CBS | rep1 | WT | yes | yes | (*19*) |
| Sox2_-116kb | pME013 | 1xCBS_Rv | CBS | rep2 | WT | yes | yes | (*19*) |
| Sox2_-116kb | CRE6 | neutral_2 | neutral sequence | rep1 | WT | yes | yes | (*19*) |
| Sox2_-116kb | pME034 | Sox2P | genomic promoter | rep1 | ΔCBS-in-SCR | yes | yes | (*19*) |
| Sox2_-116kb | pME034 | Sox2P | genomic promoter | rep1 | ΔCBS-DS-SCR | yes | yes | (*19*) |
| Sox2_-116kb | pME034 | Sox2P | genomic promoter | rep2 | WT | yes | yes | (*13*) |
| Sox2_-116kb | pME034 | Sox2P | genomic promoter | rep3 | WT | yes | yes | (*13*) |
| Sox2_-116kb | pMK01 | 3xCBS_Fw_Sox2P | CBS+promoter | rep1 | WT | yes | yes | (*19*) |
| Sox2_-116kb | pMK02 | 3xCBS_Rv_Sox2P | CBS+promoter | rep1 | WT | yes | yes | (*19*) |
| Sox2_-116kb | pMK03 | 1xCBS_Fw_Sox2P | CBS+promoter | rep1 | WT | yes | yes | (*19*) |
| Sox2_-116kb | pME034 | Sox2P | genomic promoter | rep2 | ΔCBS-in-SCR | yes | yes | (*19*) |
| Sox2_-116kb | pME034 | Sox2P | genomic promoter | rep2 | ΔCBS-DS-SCR | yes | yes | (*19*) |
| Sox2_-116kb | pME038 | 3xCBS_Fw | CBS | rep2 | WT | yes | yes | (*19*) |
| Sox2_-116kb | CRE6 | neutral_2 | neutral sequence | rep2 | WT | yes | yes | (*19*) |
| Sox2_-116kb | pMK01 | 3xCBS_Fw_Sox2P | CBS+promoter | rep2 | WT | yes | yes | (*19*) |
| Sox2_-116kb | pMK02 | 3xCBS_Rv_Sox2P | CBS+promoter | rep2 | WT | yes | yes | (*19*) |
| Sox2_-116kb | pMK03 | 1xCBS_Fw_Sox2P | CBS+promoter | rep2 | WT | yes | yes | (*19*) |
| Sox2_-116kb | pME034 | Sox2P | genomic promoter | rep4 | WT | yes | yes | (*13*) |
| Tdgf1_137 | pLD037 | neutral_1 | neutral sequence | rep1 | WT | no | no |  |
| Tdgf1_3 | pLD037 | neutral_1 | neutral sequence | rep1 | WT | no | no |  |
| Tdgf1_48 | pLD037 | neutral_1 | neutral sequence | rep1 | WT | yes | yes | (*18*) |
| Tdgf1_1 | pLD037 | neutral_1 | neutral sequence | rep1 | WT | yes | yes | (*18*) |

### Supplementary figures

**Figure S1.**
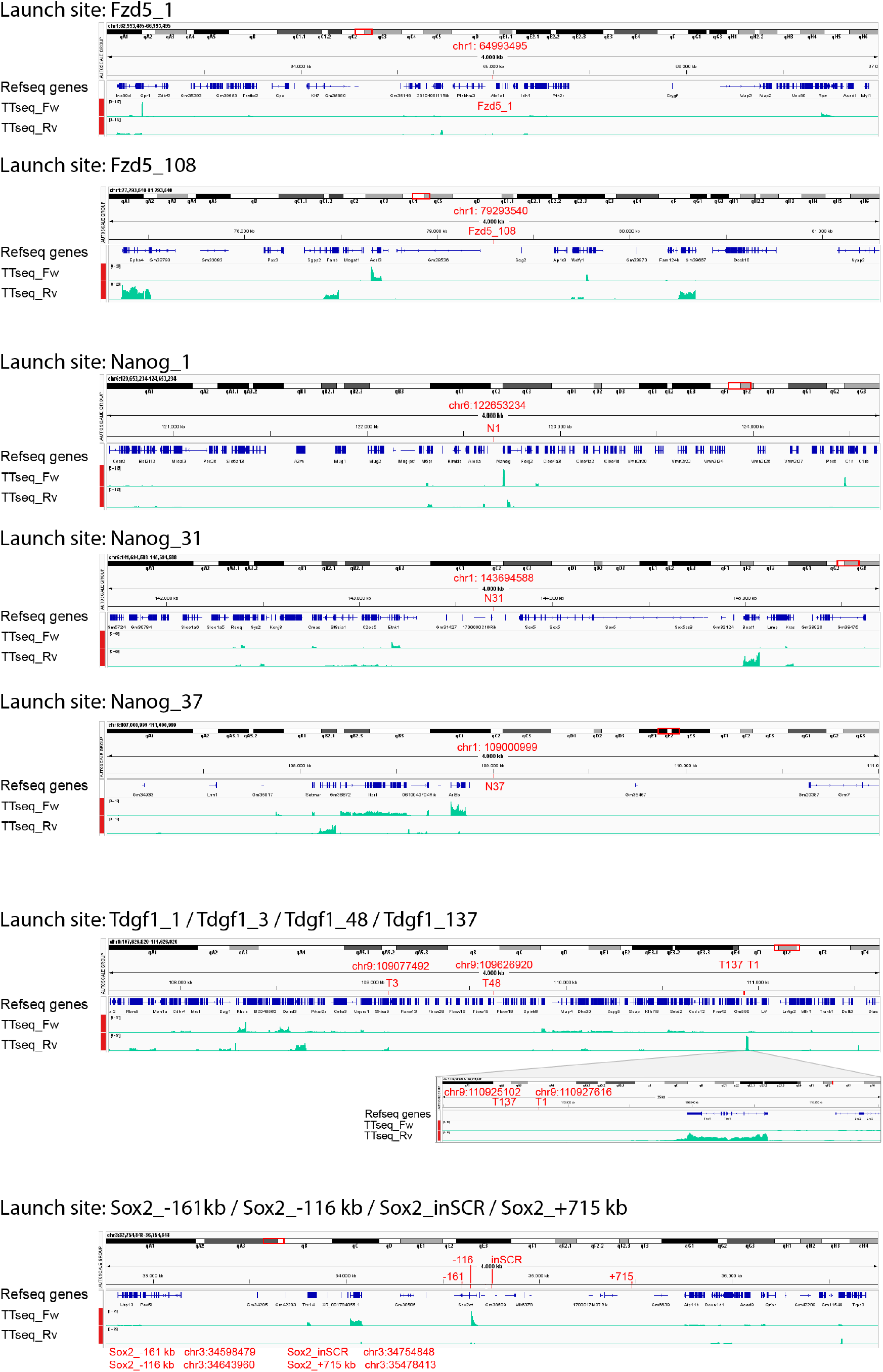
Genomic context surrounding the launch sites. IGV Genome browser snapshots showing the genomic environment of the single Sleeping Beauty launch sites used in this study. Red ticks mark the genomic position of each launch site, with the corresponding chromosome coordinate indicated in red using the mm10 genome assembly. For loci containing multiple launch sites, all positions are shown with the same genomic window. Blue tracks show RefSeq gene annotations. Green tracks show nascent transcription measured by TT-seq on the forward and reverse genomic strands (64).

**Figure S2.**
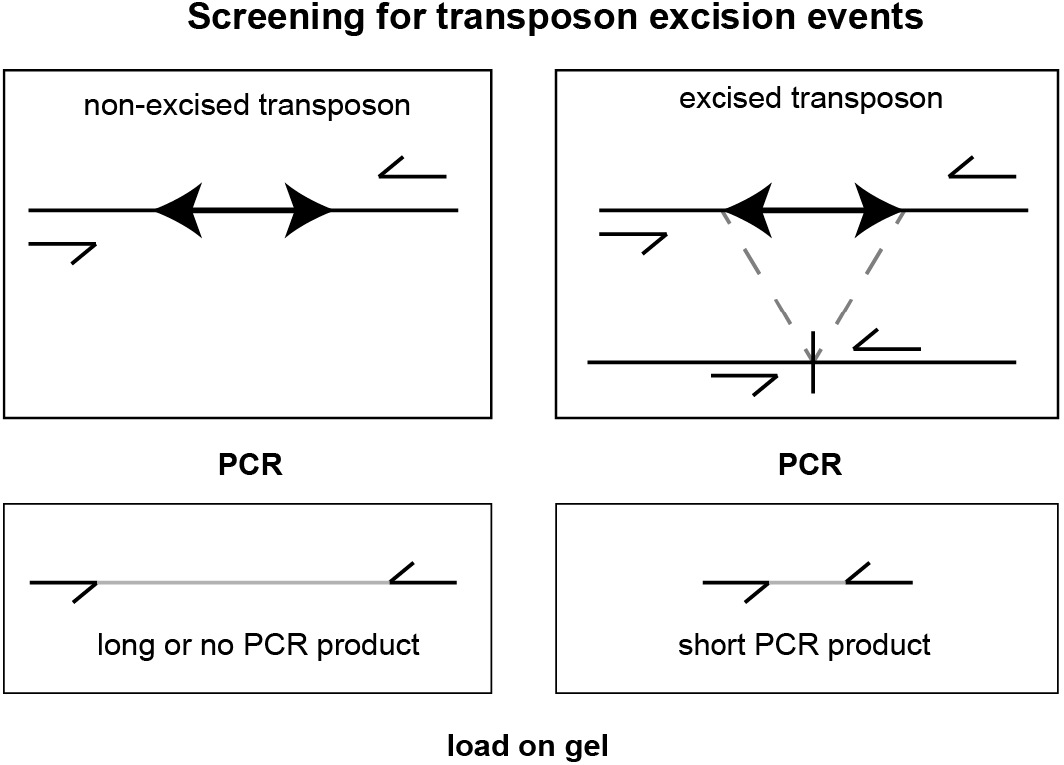
PCR screening of clones with transposon excision events. Clones were screened for successful transposon excision by PCR with primers spanning the launch site. Resulting PCR products were loaded on gel, short PCR products corresponded to excised transposon, large or no PCR product corresponded to no excision event. Size of PCR products vary per launch site.

**Figure S3.**
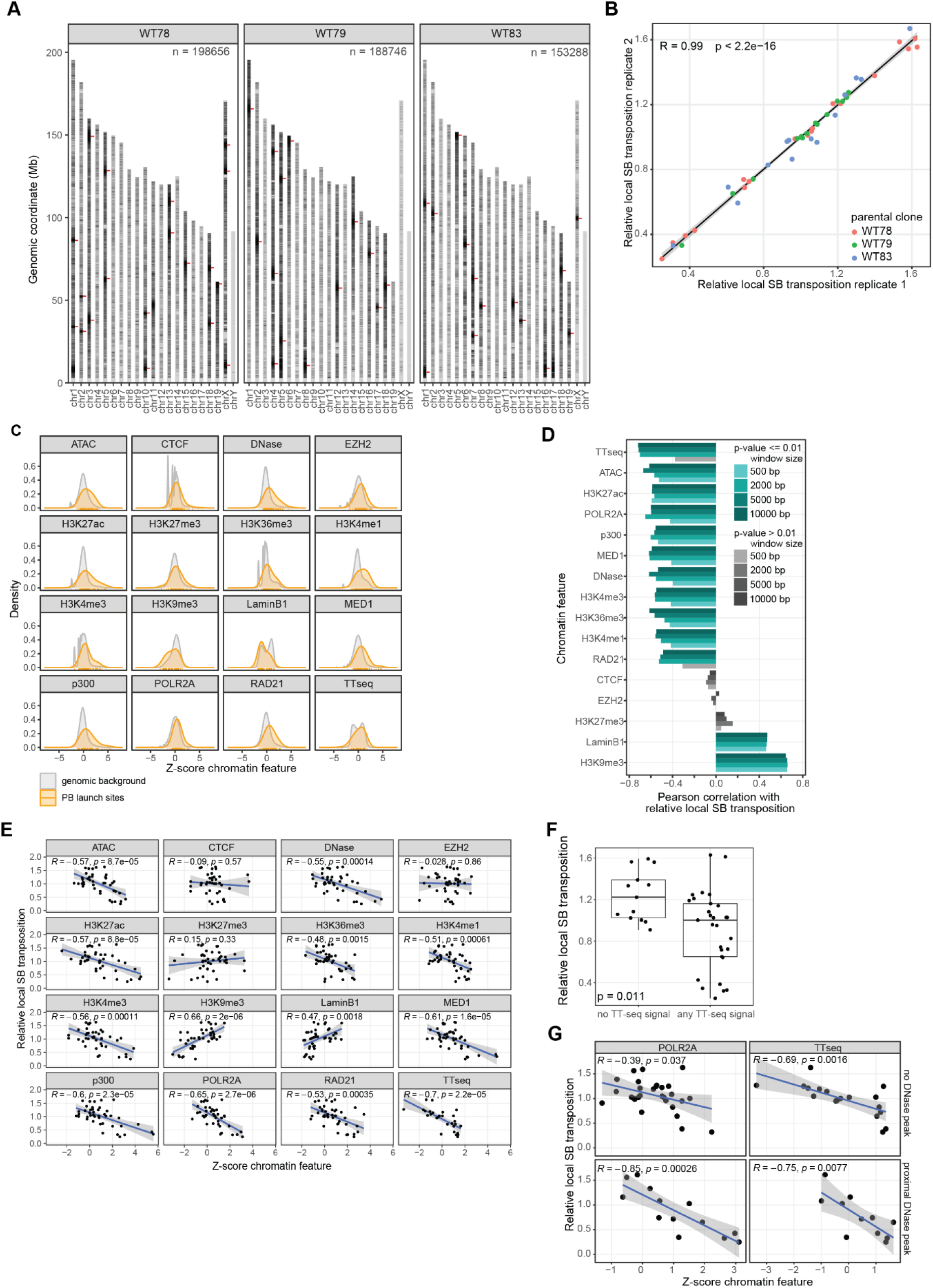
Chromatin context launch site affects transposition efficiency. **(A)** SB reintegrations mapped after SB transposase transfection in WT78. Red ticks indicate the mapped launch sites. Two biological replicates combined. n = total number of mapped SB reintegrations. WT78 also shown in **Fig 2B**. **(B)** Correlation of relative local SB transposition (number of local reintegrations, normalized to the mean per parental clone) across two biological replicates. R is Pearson correlation coefficient. **(C)** Signal of the 16 chromatin features in a 2 kb window around each of the 42 launch sites (yellow) and a background set of 879275 random 2 kb windows across the genome (grey). For the launch sites, ticks indicate the individual observations. Z-score is truncated at [-8,8]. **(D)** Pearson correlations between relative local SB transposition and each of the 16 chromatin features (ordered by correlation coefficient), across the 42 launch sites. Chromatin signal is calculated for four window sizes around the launch sites (500 – 10,000 bp). Color indicates significance (p ≤ 0.01), shade indicates window size. 2000 bp window size also shown in **Fig 2G**. **(E)** Correlation between relative local SB transposition and each of the 16 chromatin features (z-score normalized), across the 42 launch sites. R is Pearson correlation coefficient. **(F)** Relative local SB transposition for launch sites with and without any TT-seq signal. Statistical significance calculated using Wilcoxon signed-rank test. **(G)** Correlation between relative local SB transposition and POLR2A ChIP-seq or TT-seq signal (calculated in a 2 kb window), for launch sites with and without a DNase peak in the 4 kb window around the launch site. R is Pearson correlation **(C, E-G)** Chromatin signal is quantified in a 2 kb window around each launch site, and z-score normalized on a genome-wide background set. ChIP and pA-DamID data are pseudocounted. Source data provided in Methods. **(D, E, G)** For TT-seq, launch sites with no signal in the given window are excluded.

**Figure S4.**
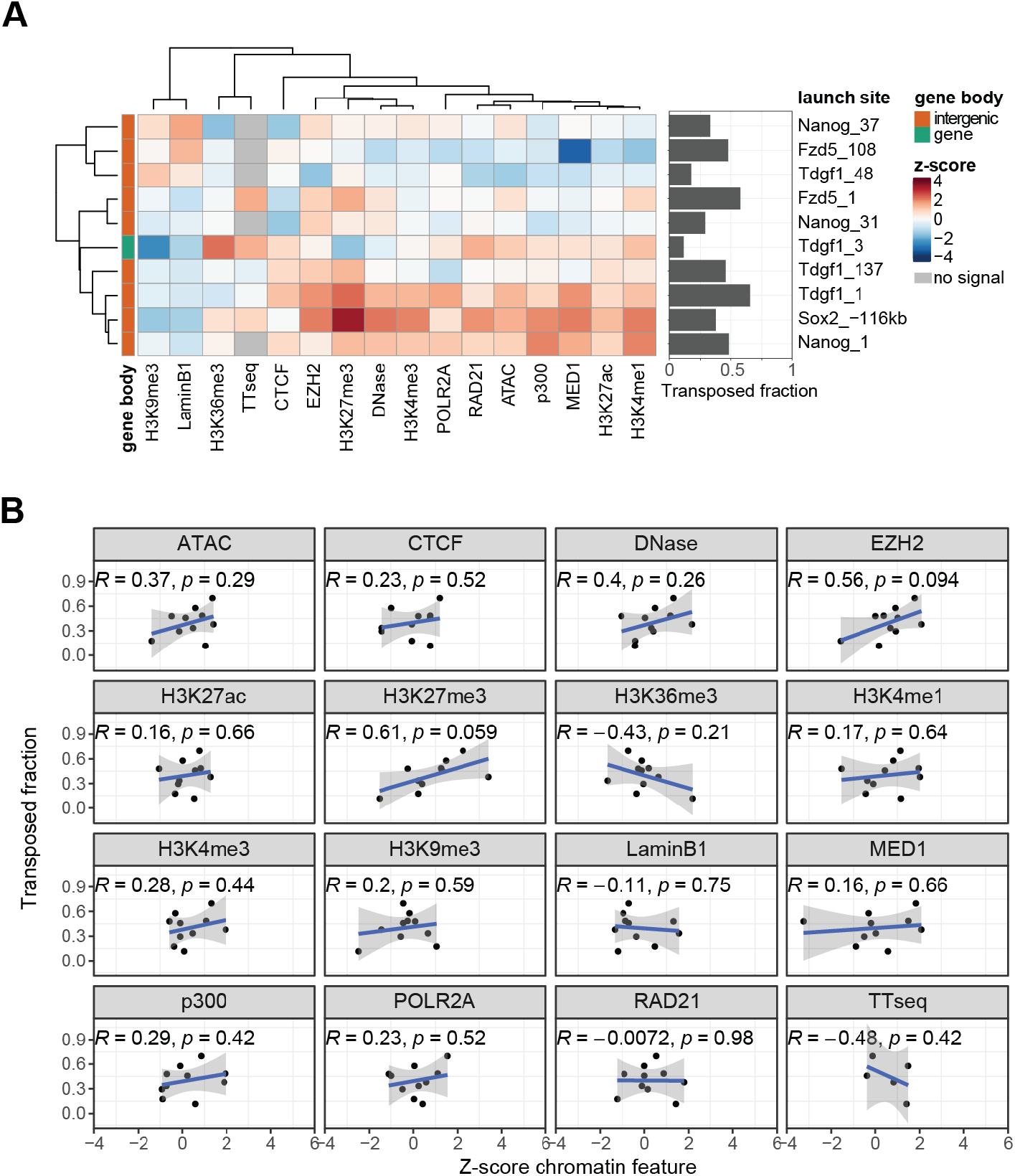
Chromatin context does not explain differences between single launch site cell lines. **(A)** Heatmap of 16 chromatin features in a 2 kb window around each of the 10 launch sites from individual launch site cell lines. Features are z-score normalized based on a genome-wide background set, for visualization the scale is truncated at [-4,4]. ChIP and pA-DamID data are pseudocounted, for TTseq no signal is indicated in grey. Source data provided in Methods. Side barplot indicates the mean transposed fraction per launch site (number of transposed reads divided by total reads). Two biological replicates for Nanog_1 and Sox2-116 kb, one for the other launch sites. Transposed fraction also shown in **Fig 1E**. **(B)** Correlation between transposed fraction and each of the 16 chromatin features (z-score normalized), across the 10 launch sites. R is Pearson correlation coefficient.

**Figure S5.**
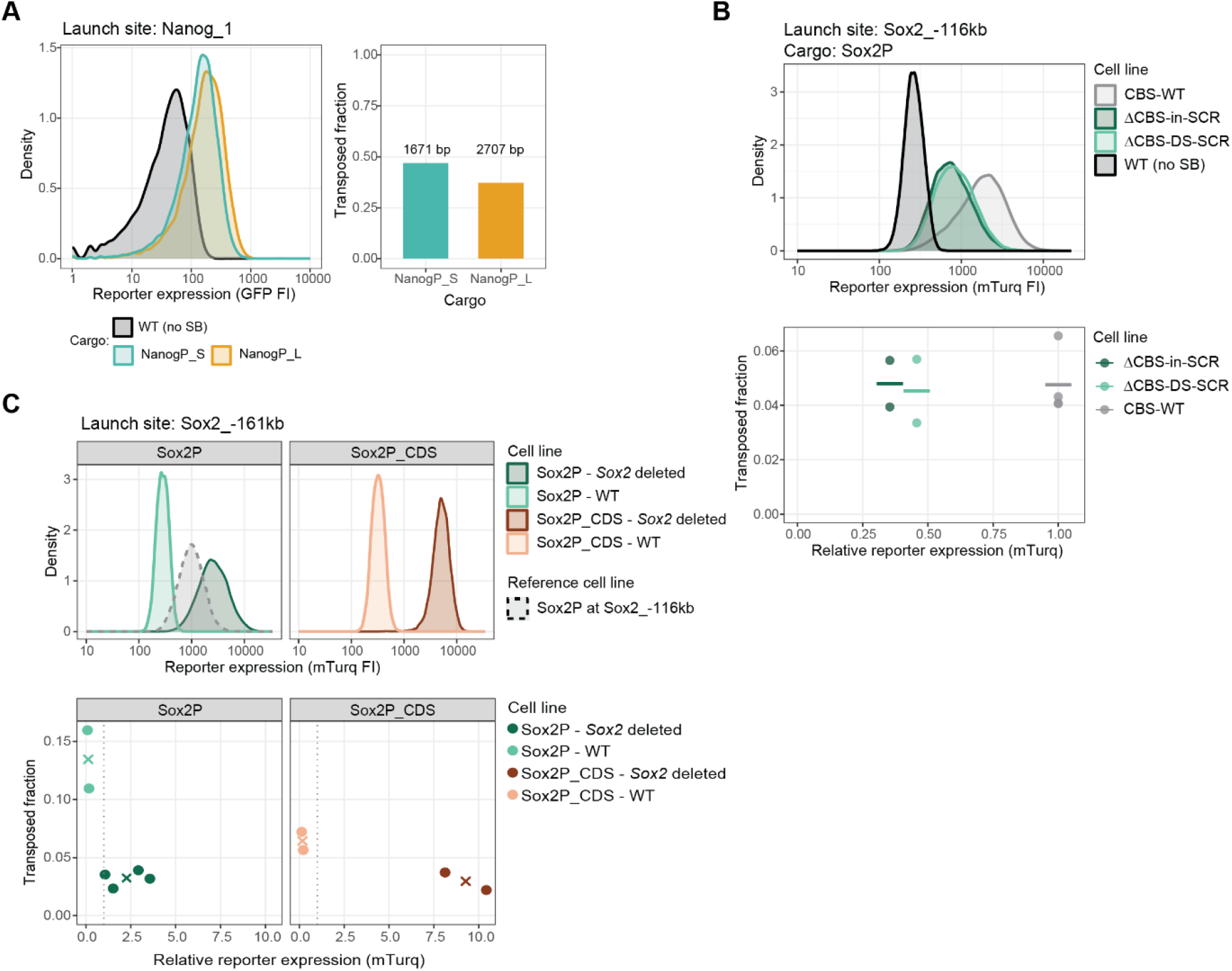
Transposition efficiency depends on cargo activity and size. **(A)** Left: Reporter expression (GFP) of two Nanog reporter cargos (NanogP_S & NanogP_L) inserted in the Nanog_1 launch site, together with a non-fluorescent control, measured by flow cytometry. For NanogP_S only the variant matched to NanogP_L is used here, see methods for details. Right: Transposed fraction of both reporter cargos and their corresponding cargo size in base pairs (bp). Colors indicated the reporter cargos. **(B)** Top: Sox2P reporter expression (mTurquoise2) from the Sox2_-116 kb launch site in different genetic backgrounds, measured by flow cytometry. In ΔCBS-in-SCR and ΔCBS-DS-SCR a CBS in or downstream (DS) of the Sox2 control region (SCR) was deleted, see see (*19*) for details. Bottom: Relationship between relative reporter expression and transposed fraction in the same cell lines. Dots indicate biological replicates colored by cell line, line indicates mean. **(C)** Top: Reporter expression (mTurquoise2) of two transposon cargos (Sox2P and Sox2P_CDS) at the Sox2_-161kb launch site in the presence (WT) or absence (Sox2 deletion) of the endogenous Sox2 gene, measured by flow cytometry. Grey density plot indicates a reference cell line carrying Sox2P at the Sox2_-116 kb launch site in the presence of the endogenous Sox2 (CBS-WT in B). Bottom: Relationship between relative reporter expression and the transposed fraction in the same cell lines. Dots indicate biological replicates colored by cell line, crosses indicate the mean per cell line. The vertical dotted line at 1 indicates reporter expression of the reference cell line. **(B-C)** Relative reporter (mTurq) expression is calculated by subtracting autofluorescence (based on a wild-type control) and normalizing to the reference cell line (Sox2P in Sox2_-116kb, without CBS or Sox2 deletion, shown in grey in B and C). **(A-C)** FI is fluorescent intensity.

**Figure S6.**
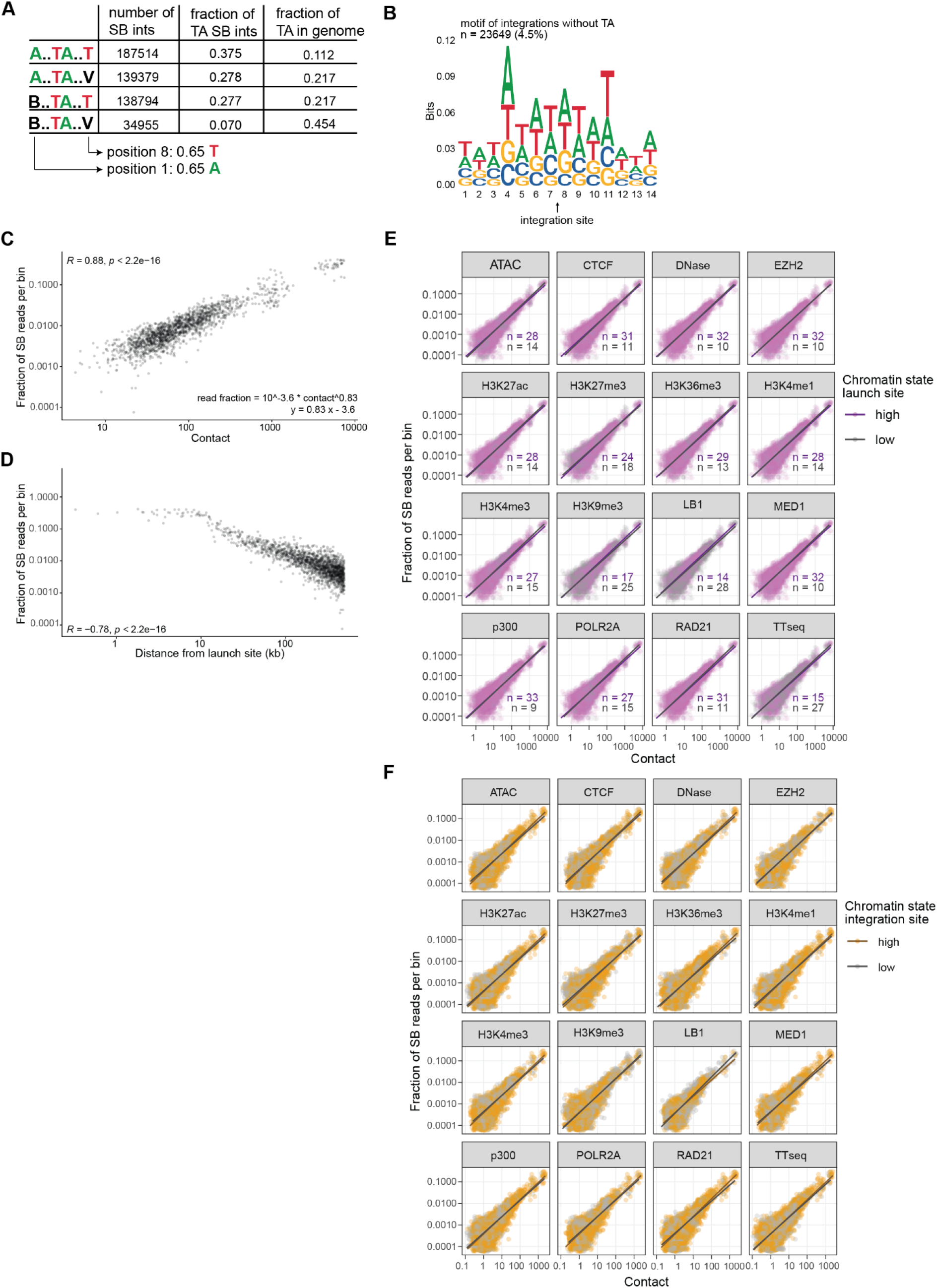
3D contact determines reintegration distribution. **(A)** Number of SB reintegrations found at each of the four possible sequence motifs around a TA site. Columns indicate: number of SB reintegration sites matching each pattern; fraction of all SB reintegrations at TA sites matching each pattern; fraction of total TA sites in the genome matching each pattern. **(B)** Sequence motif derived from the 23649 mapped SB reintegration sites not a a TA site (4.5% of all mapped reintegrations). **(C)** Correlation between the fraction of SB mapping reads and the contact with the launch site per 25 kb bin within 500 kb of each of the 52 launch sites. **(D)** As in C, but for the relationship between the genomic distance from the launch site and the fraction of SB mapping reads, calculated on the same 25 kb bins. **(C-D)** R is Spearman correlation coefficient, data is the same as in Fig 4D-E but restricted to bins within 500kb of the launch site. **(E)** Relationship between the fraction of SB mapping reads per 25 kb bin within 2 Mb of a launch site and contact frequency of that bin with the launch site, stratified by high (z-score > 0) or low (z-score ≤ 0) levels of various chromatin features at the launch site. Chromatin level is calculated in a 2 kb bin surrounding the launch site (as in Fig 2F). n indicates the number of launch sites in each group. H3K9me3 is also shown in Fig 4G**. (F)** Relationship between the fraction of SB mapping reads per 10 kb bin within 2 Mb of a launch site and contact frequency of that bin with the launch site, stratified by high (z-score > 0) or low (z-score ≤ 0) levels of various chromatin features at the 10 kb reintegration bin. Launch sites with less than 30% high or low bins are excluded, n indicates the number of launch sites included per feature. ATAC is also shown in Fig 4I. (C-F) Bins with 0 contact or 0 reintegrations are excluded. **(A-F)** SB reintegration data from the nested transposon experiment described in Fig 2. Data pooled from three cell lines, two biological replicates per cell line.

**Figure S7.**
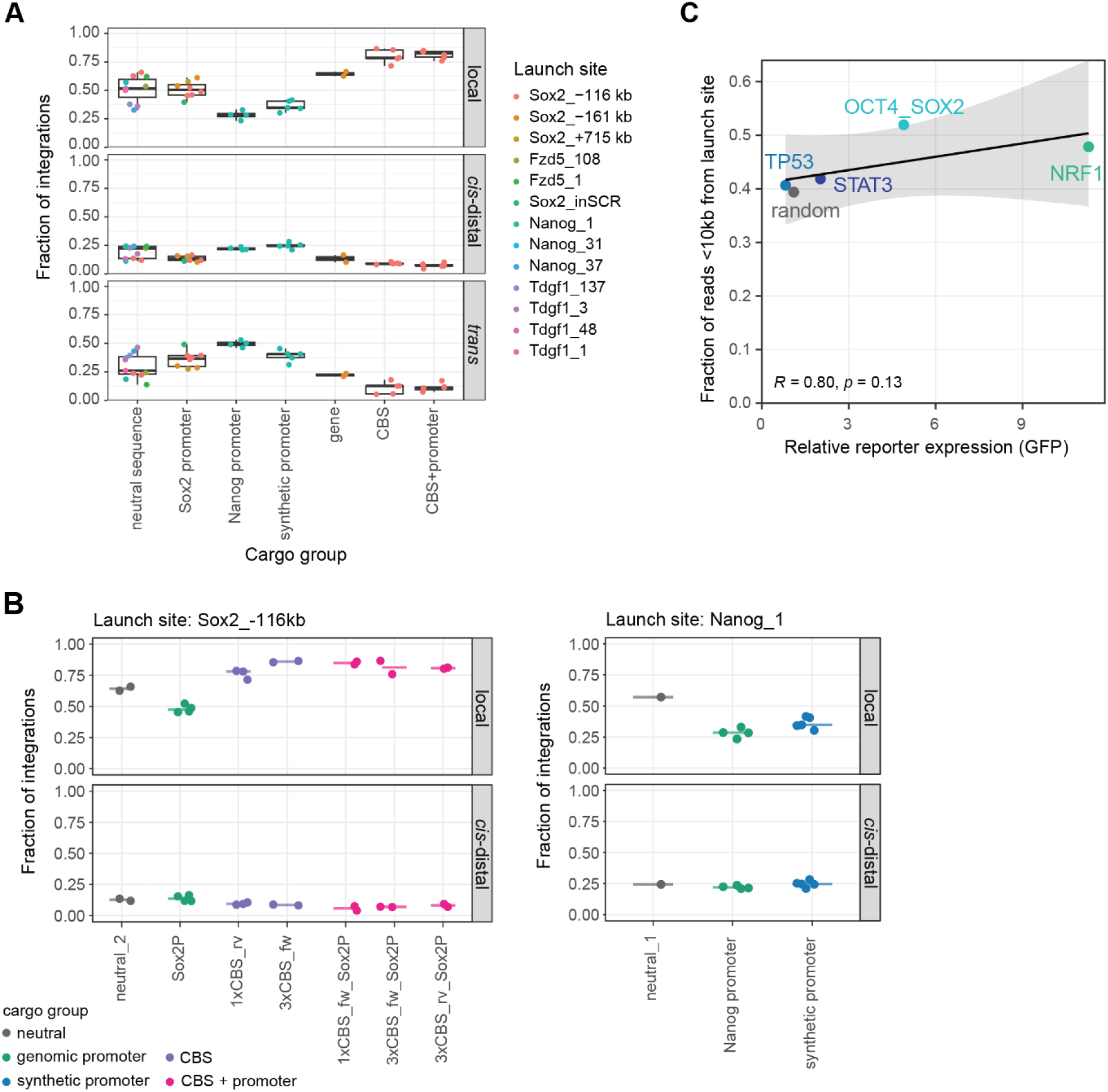
Cargo characteristics shape local and interchromosomal reintegration patterns. **(A)** Fraction of reintegrations classified as local (within 2 Mb of the launch site), *cis*-distal (same chromosome but >2 Mb from the launch site) or *trans* (other chromosomes) per cargo group. Dots represent biological replicates and are coloured by launch site. **(B)** Fraction of reintegrations classified as local and *cis*-distal for different cargos mobilized from the Sox2_-116 kb and Nanog_1 launch sites. Dots represent biological replicates and horizontal lines indicate the median. **(C)** Relationship between the reporter expression and the fraction of all local (<2 Mb) SB mapping reads found within 10 kb of the launch site, for the five synthetic promoters mobilized from launch site Nanog_1. Relative reporter expression (GFP) is calculated by subtracting autofluorescence (based on a wild-type control) and normalizing to the random reporter. R is Spearman correlation coefficient, colour indicates the synthetic promoter identity, one biological replicate.

